# Crowding changes appearance systematically in peripheral, amblyopic, and developing vision

**DOI:** 10.1101/2021.11.30.470647

**Authors:** A.V. Kalpadakis-Smith, V.K. Tailor, A.H. Dahlmann-Noor, J.A. Greenwood

## Abstract

Visual crowding is the disruptive effect of clutter on object recognition. Although most prominent in adult peripheral vision, crowding also disrupts foveal vision in typically-developing children and those with strabismic amblyopia. Do these crowding effects share the same mechanism? Here we exploit observations that crowded errors in peripheral vision are not random: target objects appear either averaged with the flankers (assimilation), or replaced by them (substitution). If amblyopic and developmental crowding share the same mechanism then their errors should be similarly systematic. We tested foveal vision in children aged 3-8 years with typical vision or strabismic amblyopia, and peripheral vision in typical adults. The perceptual effects of crowding were measured by requiring observers to adjust a reference stimulus to match the perceived orientation of a target ‘Vac-Man’ element. When the target was surrounded by flankers that differed by ±30°, all three groups (adults and children with typical or amblyopic vision) reported orientations between the target and flankers (assimilation). Errors were reduced with ±90° differences, but primarily matched the flanker orientation (substitution) when they did occur. A population pooling model of crowding successfully simulated this pattern of errors in all three groups. We conclude that the perceptual effects of amblyopic and developing crowding are systematic and resemble the near periphery in adults, suggesting a common underlying mechanism.

**Precis:** Crowding strongly disrupts peripheral vision, as well as the foveal vision of children with typical vision and amblyopia. We show that typically developing and amblyopic children make the same crowded errors as adults in the visual periphery, consistent with a common mechanism in all three cases.

## Introduction

Clutter can significantly disrupt the recognition of objects that are otherwise readily identified in isolation – a phenomenon known as visual crowding (Levi, 2008; Whitney & Levi, 2011). In the typical adult visual system, crowding is most pronounced in peripheral vision, where the recognition of a target can be hindered by flanking objects separated by as much as half the target eccentricity (Bouma, 1970). Although this disruption is usually minimal in the fovea (Toet & Levi, 1992; Danilova & Bondarko, 2007; Coates, Levi, Touch, & Sabesan, 2018), elevations occur in amblyopia, a developmental disorder of vision characterised by reduced acuity in one eye despite optical correction (McKee, Levi, & Movshon, 2003). When amblyopia is associated with strabismus (ocular misalignment), foveal vision in the affected eye is strongly impaired by the presence of nearby flankers in both adults (Flom, Weymouth, & Kahneman, 1963; Levi & Klein, 1985) and children (Greenwood et al., 2012). Foveal elevations in crowding have also been found in children with typical vision up to the age of 11 years (Jeon, Hamid, Mauer, & Lewis, 2010; Greenwood et al., 2012). These elevations have a range of functional consequences, given for instance the correlation between crowding and reading ability (Martelli, Di Filippo, Spinelli, & Zoccolotti, 2009). However, although it is clear that crowding occurs in these three instances – typical peripheral vision, the amblyopic fovea, and the developing fovea – it is unclear whether the underlying mechanism is the same.

In the typical adult periphery, crowding strongly disrupts target identification when flankers are near the target. The spatial extent of crowding can be defined as the target-flanker separation required to remove this disruptive effect (Bouma, 1970; Toet & Levi, 1992). These values are typically far larger than what would be predicted by the level of acuity or blur in peripheral vision (Song, Levi, & Pelli, 2014) and do not vary with target size (Levi, Hariharan, & Klein, 2002b; Tripathy & Cavanagh, 2002; Pelli, Palomares, & Majaj, 2004). Peripheral crowding is also selective for the similarity between the target and flankers in visual dimensions such as orientation and contrast polarity, with target identification more strongly disrupted when flankers are similar to the target than when they are dissimilar (Kooi, Toet, Tripathy, & Levi, 1994; Wilkinson, Wilson, & Ellemberg, 1997; Chung, Levi, & Legge, 2001). In the amblyopic fovea, crowding is similarly reduced by an increase in target-flanker separation (Levi, Hariharan, & Klein, 2002a; Hariharan, Levi, & Klein, 2005), with an extent that is also well in excess of limits imposed by acuity or blur (Song, Levi, & Pelli, 2014) and largely invariant to target size (Levi, Hariharan, & Klein, 2002a). Amblyopic crowding may however be less dependent on target-flanker similarity – flankers dissimilar to the target in polarity, contrast, and orientation have been found to be equally disruptive as flankers similar to the target along these dimensions (Levi, Hariharan, & Klein, 2002a; Hariharan, Levi, & Klein, 2005). These variations in the characteristics of crowding cast doubt on the possibility of a common mechanism for peripheral vision and the amblyopic fovea. Although crowding in the typically developing fovea is also dependent on target-flanker separation (Greenwood et al., 2012), the effects of acuity/blur, target size, and target-flanker similarity are unclear.

Much of our understanding of the mechanisms underlying crowding in the typical adult periphery derives from measurements of the errors that observers make when reporting the identity of a crowded target. Observers have been found to report either the identity of one of the flankers surrounding the target (Strasburger, Harvey, & Rentschler, 1991; Strasburger, 2005), or an intermediate identity close to the target-flanker average (Parkes et al., 2001; Greenwood, Bex, & Dakin, 2009). The finding that target patches of noise can similarly adopt the perceived orientation of the flankers (Greenwood, Bex, & Dakin, 2010) suggests that these errors are not simply the result of decisional bias. Rather, they represent a change in the perceived identity of the target to more closely resemble the flankers, indicating that peripheral crowding has systematic perceptual effects.

A range of models can account for this systematic shift in the identity of crowded targets. Substitution models (Ester, Klee, & Awh, 2014; Ester, Zilber, & Serences, 2015) argue that errors emerge due to the substitution of a flanker into the target location, leading observers to report the flanker identity. This substitution is either attributed to the increased positional uncertainty of peripheral vision (Wolford, 1975; Krumhansl & Thomas, 1977), or unfocussed spatial attention (Strasburger, Harvey, & Rentschler, 1991; Strasburger, 2005). On the other hand, ‘pooling’ or averaging models (Parkes et al., 2001; Greenwood, Bex, & Dakin, 2009; Dakin, Cass, Greenwood, & Bex, 2010) posit that crowding is the compulsory integration of target and flanker signals, resulting in observers perceiving an average or intermediate feature (e.g. orientation) of the target and flankers. Each of these model types focuses on distinct types of errors: either flanker reports (‘substitution errors’) or reports of intermediate identities between the target and flankers (‘assimilation errors’).

Population pooling models (van den Berg, Roerdink, & Cornelissen, 2010; Harrison & Bex, 2015; Greenwood & Parsons, 2020) propose a more general framework for the perceptual effects of peripheral crowding. Harrison and Bex (2015) used an orientation-matching task where observers matched the orientation of a reference Landolt-C to that of a crowded target in the periphery. When the target was surrounded by flankers that differed by 45° or less, observers reported orientations between the target and flanker values (assimilation errors). When the difference from the target was 90° or above, errors more closely matched the flanker orientation (substitution errors). Rather than invoking separate substitution or averaging mechanisms, Harrison and Bex (2015) account for both error types using a population pooling model that takes a weighted combination of population responses to the target and flankers. Similar approaches have been applied more generally to explain crowding with letters (Freeman, Chakravarthi, & Pelli, 2012) and faces (Kalpadakis-Smith, Goffaux, & Greenwood, 2018). Higher-dimensional pooling approaches have also been developed, which depict crowding as an over-application of summary statistics across the periphery (Balas, Nakano, & Rosenholtz, 2009; Freeman & Simoncelli, 2011; Keshvari & Rosenholtz, 2016). Their generality allows the consideration of these crowding effects in a range of naturalistic tasks (Rosenholtz, Yu, & Keshvari, 2019), though quantitative predictions of these high-dimensional models are more difficult to discern for specific paradigms.

Although population pooling models can account for peripheral crowding, their applicability to amblyopia is unknown. Given the plethora of deficits in visual function observed in the affected eye (McKee, Levi, & Movshon, 2003), the basis of amblyopic crowding could in fact differ substantially from that of peripheral vision. In addition to the definitive acuity deficit, vision in the affected eye of observers with amblyopia is characterised by increased positional uncertainty (Levi & Klein, 1985). This uncertainty could produce confusions of the flanker for the target, making a predominance of substitution errors. Alternatively, crowded errors may arise due to perceptual distortions that affect the amblyopic eye. Observers with strabismic amblyopia show considerable distortions when reproducing visual stimuli, including shrinkage, expansion, and torsion of specific regions (Pugh, 1958; Sireteanu, Lagreze, & Constantinescu, 1993). Although these distortions are consistent over time, they vary across observers and visual field location (Barrett et al., 2003). If these distortions underlie the perceptual effects of amblyopic crowding, then errors would not be systematic but random (depending on the particular conjunction of the distortion type and stimulus), suggesting a distinct mechanism from peripheral crowding.

Even less is known about the mechanism of foveal crowding during development. Although the extent of foveal crowding has been found to be greater in typically developing children than in adults (Atkinson & Braddick, 1983; Atkinson, Anker, Evans, & McIntyre, 1987; Jeon et al., 2010; Greenwood et al., 2012), the perceptual effects of developing crowding have not been investigated. Studies have however demonstrated that children make a disproportionate amount of random errors in psychophysical tasks relative to adults (Witton, Talcott, & Henning, 2017; Manning, Jones, Dekker, & Pellicano, 2018). These errors are frequently made in low difficulty “catch” trials (Treutwein, 1995) and have been attributed to attentional lapses and under-developed short-term memory. Both of these factors would produce random errors that could dominate responses to the identity of a target object in crowding paradigms. The same could be true of children with amblyopia. In both cases, the observed elevations in foveal crowding could therefore reflect quite distinct processes to the systematic perceptual effects observed in peripheral vision.

To investigate the perceptual effects of amblyopic and developing crowding, we tested the foveal vision of children aged 3-8 years with either strabismic amblyopia or typical vision, with comparison to peripheral vision in typical adults. Though many studies of amblyopic crowding have used adult observers, we felt it important to test these deficits in children, given that both the onset and treatment of this condition typically occurs in early childhood (Holmes et al., 2011; Tailor, Bossi, Greenwood, & Dahlmann-Noor, 2016). To this end, we adapted the orientation-matching task used by Harrison and Bex (2015), with stimuli presented foveally to children with and without amblyopia, and in peripheral vision to adults. Given the timing constraints in testing children (due to shorter attention spans), we tested ±30° and ±90° target-flanker orientation differences. The ±30° flanker differences were selected to test whether children show the same systematic shift of responses to orientations between the target and flankers (assimilation errors) as in the periphery. Because orientation differences below 90° are less able to distinguish between assimilation and substitution errors (Harrison & Bex, 2015), we chose ±90° flanker differences to further constrain the underlying mechanisms. These ±90° orientation differences have also been shown to reduce the effect of crowding in the periphery relative to smaller orientation differences (Hariharan, Levi, & Klein, 2005; Harrison & Bex, 2015), allowing us to examine the selectivity for target-flanker similarity in amblyopic and developing crowding. We further probed the possibility of a common mechanism by simulating the observed perceptual effects with a population pooling model of crowding, with comparison to a model that simply added noise to the orientation judgements.

If there is a common mechanism that underlies amblyopic, developing, and peripheral crowding, each of these instances should show the same systematic effects on target appearance as observed in the adult periphery. That is, children with amblyopia and typical vision should make either assimilation or substitution errors, depending on the target-flanker orientation difference. In contrast, distinct mechanisms for crowding in strabismic amblyopia and developing vision may produce random errors, either because crowding affects the appearance of the target in a non-systematic manner or due to attentional lapses.

## Methods

### Design

Both children and adults completed three tasks. Acuity was measured first to determine the minimum target size at which each observer could judge the orientation of the target element. Crowded acuity was then measured to determine the extent of the spatial zone of crowding around the target location. These tasks, adapted from Greenwood et al. (2012), were used to set stimulus sizes for the orientation-matching task, in particular to ensure both that the target size was above acuity limits and that flankers were within the spatial region of crowding. The third orientation-matching task, adapted from Harrison & Bex (2015), allowed us to measure the perceptual effects of crowding.

### Observers

#### Children

40 children between 3-8 years of age were tested, divided into two groups: those with typical vision (*n*= 20, mean= 73.2 months, range= 36-94 months), and those with strabismic amblyopia (*n*= 20, mean= 70.7 months, range= 37-100 months). Sample sizes were derived from prior work (Greenwood et al., 2012), with children tested at the Children’s Eye Centre at Moorfields Eye Hospital (London, UK).

Prior to the study, children underwent a full orthoptic assessment to ensure they met our inclusion and exclusion criteria. Typically developing children were selected to have a best-corrected visual acuity of 0.1 logMAR (logarithm of the minimum angle of resolution) or better in both eyes, as measured by Thomson V2000 acuity charts, in the absence of any pre-existing visual or neurological deficits. For children with strabismic amblyopia, inclusion was based on the presence of amblyopia, as indicated by a two-line difference in best-corrected logMAR acuity between the eyes, as well as heterotropia (deviation of the optical axes) that could be either esotropia (inward deviation) or exotropia (outward). Children with additional visual deficits (e.g. macular dystrophies) and developmental or neurological conditions (e.g. autism) were excluded. We did not exclude cases of joint anisometropia and strabismus. Clinical details for all children are shown in Appendix A.

Three children with amblyopia did not complete all experimental tasks and were excluded from the analysis. They are not included in the tallies above. The experimental procedures were performed with the informed consent of children and their parents and were approved by the Health Research Authority of the UK National Health Service.

#### Adults

10 adults were tested (4 males, mean= 28.7 years, range= 24-35 years), including two of the authors (AKS and JAG). All had a best-corrected visual acuity of 0 logMAR or better. None had amblyopia or strabismus, nor any history of binocular dysfunction, as indicated by self-report.

### Apparatus

#### Children

Experiments were programmed using MATLAB (The Mathworks, Ltd., Cambridge, UK) and run on a Dell PC using PsychToolbox (Brainard, 1997; Pelli, 1997). Stimuli were presented on an ASUS VG278HE LCD monitor, with 1920 × 1080 resolution and 120 Hz refresh rate. The monitor was calibrated using a Minolta photometer and linearised in software, to give a maximum luminance of 150 cd/m^2^. A second Dell UltraSharp 2208WFP LCD monitor, with 1680 × 1050 resolution and 75 Hz refresh rate, was positioned above the first. In the acuity and crowding-extent tasks, this second monitor was used to display a running tally of points children received by playing the games. In the orientation-matching task it displayed the response stimulus.

Figure 1A shows the experimental setup for the children. Children wore stereo-shutter glasses (nVidia Corp., Santa Clara, CA) alternating at 120 Hz, custom-fit into a ski-mask frame to allow a comfortable fit over their optical correction. These glasses were used to present the stimuli monocularly. Children were seated 3 m from the screen. For the acuity and crowding-extent tasks, the experimenter recorded the children’s responses using the keyboard. For the orientation-matching task, a Griffin Powermate response dial was used by the children to rotate the response element and register their responses.

**Figure 1.**
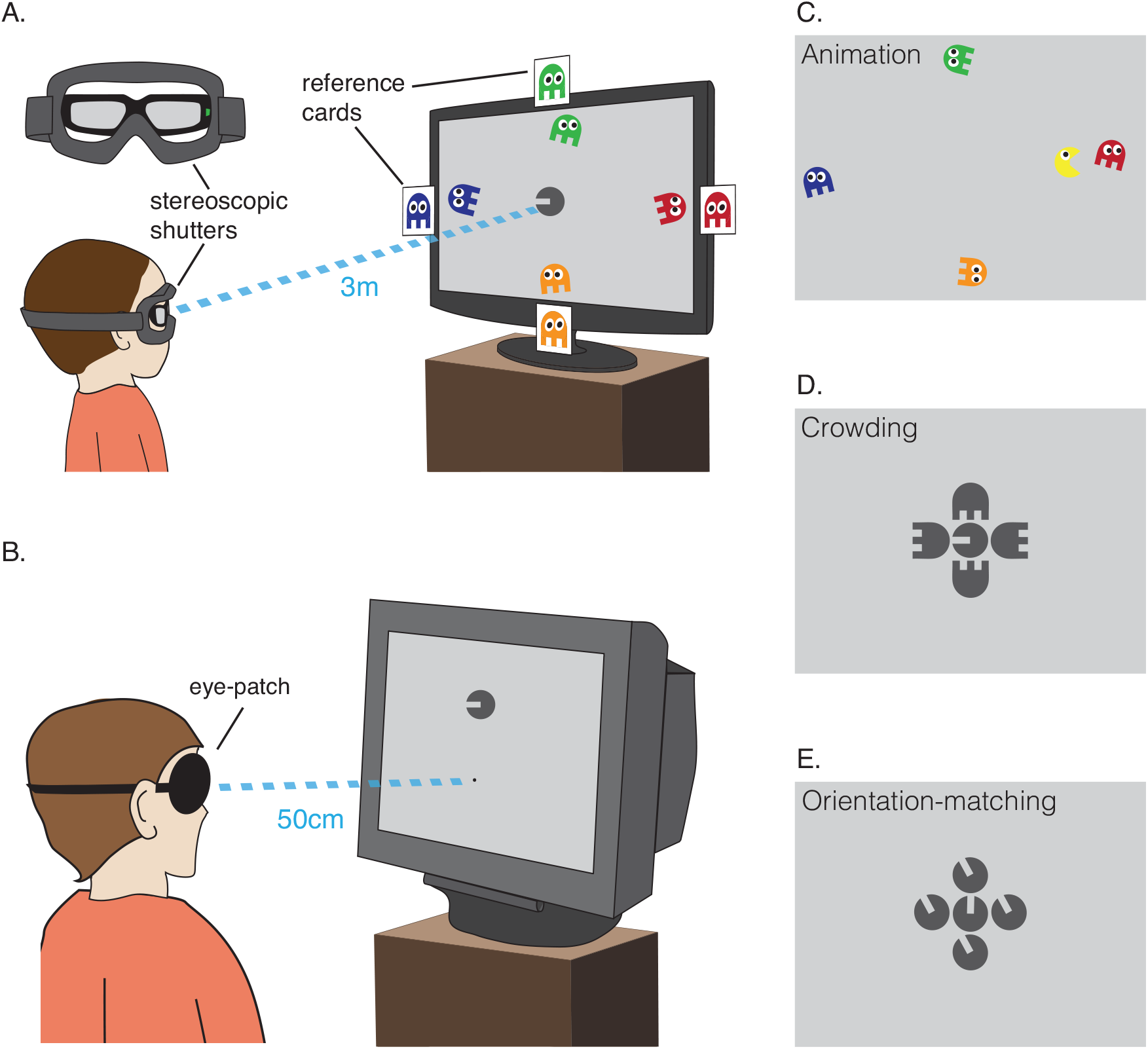
Apparatus and stimuli. **A**. For children, stimuli were viewed through stereoscopic shutter glasses mounted in a ski mask and presented on a 3D-compatible monitor at 3 m distance. An example trial of the acuity task is depicted, where children reported the colour of the ghost that Vac-Man was facing. Coloured cards of the ghosts on the monitor edges helped children select the ghost. **B**. Adults viewed the monitor from 50 cm, wearing an eye patch over their non-dominant eye. **C**. An example frame from the “reward animation”, presented every three correct trials. **D**. Illustration of the stimuli in the crowding-extent task. Ghost flankers were presented at random orientations at a fixed relative separation (1.1× stimulus diameter), with their absolute separation varied by QUEST. **E**. Illustration of stimuli in the orientation matching task. Here, flankers were filled-in Landolt-Cs, similar to the target, presented with the same orientation difference at a fixed separation.

#### Adults

Adults completed the same three tasks as children, run on a Viglen PC and presented on a Sony GDM-FW900 CRT monitor with 2304 × 1440 resolution and 80 Hz refresh rate. The monitor was calibrated and linearised to give a maximum luminance of 122 cd/m^2^. For the acuity and crowding-extent tasks, observers registered their response using a keyboard. Responses in the orientation-matching task were made with a Griffin Powermate dial. Observers were seated 50 cm from the monitor, with a head-and-chin rest used to minimise movement. Stimuli were presented monocularly to the dominant eye, with an eye-patch covering the non-dominant eye. Eye-dominance was established using the Miles test (Miles, 1928). Figure 1B depicts this experimental setup.

### Stimuli and Procedures

#### Children

The three tasks involved five video-game characters adapted from a previous study (Greenwood et al., 2012): Vac-Man (Visual Acuity Man) and four ghosts. Vac-Man was a circle with a horizontal gap for a “mouth”, resembling a filled-in Landolt-C. Prior to rotation, the mouth extended from the stimulus midpoint to the rightwards edge, with a vertical width (which we refer to as the “gap size”) that was one-fifth of the stimulus diameter, similar to the proportions in Sloan letters (Sloan, 1959). Vac-Man was the centrally located target stimulus in all three tasks, viewed foveally and rendered in black at 90% Weber contrast against a mid-grey (45 cd/m^2^) background. Vac-Man also served as flanker and response stimuli in the orientation-matching task. The ghost characters acted either as colour aids for the identification of Vac-Man’s orientation in the acuity task (as in Figure 1A), or achromatic flanker stimuli in the crowding-extent task (Figure 1D). The gap for each of the ghosts’ “legs” was also one-fifth of the stimulus diameter.

All children began with the acuity task, where they were asked to report which of the ghosts Vac-Man was facing (4 Alternative Forced Choice, 4AFC). Each ghost had a distinct colour (green above, red to the right, orange below, and blue to the left) and moved slowly along the monitor edges at a large separation from the target (see Figure 1A) to minimise the chance of any crowding with the target. Children could report either the colour of the ghost or its location, verbally or by pointing. Pictures of the ghosts were placed at the monitor edges to aid children’s reports. Normal colour-naming abilities were checked using the stimuli prior to participation. Feedback was given after each trial through brief animations, with Vac-Man smiling for correct responses and frowning when incorrect. A longer “reward” animation was presented after 3 correct responses in which Vac-Man ate a ghost (see Figure 1C). Children had unlimited response time.

In the subsequent crowding-extent task, the four ghosts became flankers surrounding Vac-Man, each achromatic in order to match the target and increase the strength of crowding through target-flanker similarity (Kooi et al., 1994). Flanker ghosts were located above, below, left, and right of Vac-Man, with each ghost randomly oriented in 1 of 4 cardinal orientations. Children made the same 4AFC judgement as the acuity task, aided by the reference cards of the ghosts on the monitor edges.

Acuity thresholds were measured by varying the overall size of Vac-Man, and thus the visibility of the mouth gap to indicate its orientation. Size was varied using a QUEST staircase procedure (Watson & Pelli, 1983) set to converge at 62.5% correct performance. These gap-size thresholds set the lower bound for the size used in the orientation-matching task. The spatial extent of crowding was also measured by varying Vac-Man size, with QUEST converging at a higher level of 80% correct performance. Flanker ghosts were scaled similarly, with the centre-to-centre separation between the target and flankers scaled at 1.1× target diameter, a value recommended as efficient for the measurement of crowding extent (Song, Levi, & Pelli, 2014). Although this method confounds size and separation by varying both, the extent of amblyopic and peripheral crowding are limited by centre-to-centre separation and not target size (Hariharan, Levi, & Klein, 2005). As such, it is only the variations in centre-to-centre separation that should affect the strength of crowding, and thus the measurement of its extent (Levi, Song, & Pelli, 2007; Song, Levi, & Pelli, 2014; Coates, Ludowici, & Chung, 2021). The resulting threshold gave the upper bound for the sizes and target-flanker separations used in the orientation-matching task, ensuring that stimuli were placed within the spatial extent of crowding.

The QUEST routine used for both acuity and crowding-extent tasks was tailored to suit children in three ways. First, to begin the task children were given 3 practice trials with a target gap-size set at twice their acuity level measured during orthoptic testing. Second, easier trials were presented on every fifth trial by selecting a gap size at twice the current QUEST threshold estimate. This minimised the frustration that numerous trials near threshold can produce. Third, an exit criterion was used to reduce testing time: if the standard deviation of the threshold estimated by QUEST for the preceding 8 trials was below 0.03 log units, the experimenter was given the option to exit the task. Otherwise, the experiment terminated after 60 trials (30 for each eye). The average number of trials needed to estimate threshold, excluding practice trials, was 44 for acuity and 46 for the crowding-extent task. Both eyes were tested in one experimental run, with separate QUEST staircases for each eye running simultaneously and selected at random on each trial. The output of each QUEST staircase gave the threshold size of Vac-Man’s mouth in degrees of visual angle.

The final orientation-matching task measured the perceptual effects of crowding. Four achromatic “imposter” Vac-Men surrounded the “real” target Vac-Man in each cardinal direction (see Figure 1E). Both target and flanker Vac-Men were at 90% Weber contrast. Here, a second response Vac-Man was presented on the response screen, twice the size of the target to ensure visibility. Children used the response dial to rotate the Vac-Man on the response screen until it appeared the same as the real Vac-Man on the main monitor. They had unlimited time to respond. On each trial, the orientation of the target varied randomly between ±45° from vertical. The four flankers were matched in orientation, all of them differing from the target by either ±30° or ±90°. This resulted in five flanker conditions: unflanked (target in isolation) or surrounded by flankers with a difference of either +30° (counter-clockwise), -30° (clockwise), +90°, or -90° from the target. 12 trials were tested for each condition, resulting in 60 trials in total for the orientation-matching task. When children’s responses deviated from the orientation of the target by more than ±35°, they received feedback in the form of a frowning Vac-Man, whereas when they responded within that range, Vac-Man smiled. This was done to maintain children’s engagement in the task and reward them for participating.

Stimulus sizes in the matching task were determined individually for each child, both to ensure that Vac-Man was visible (i.e. above the acuity limit) and crowded (i.e. with flankers within the spatial extent of crowding). A multiple of the gap-size acuity threshold was thus used, constrained by the crowding-extent values. We aimed to present stimuli with a gap size of 3× the acuity threshold, though lower values were used where this gave target-flanker separations that exceeded the crowding extent for that child (either 2.5, 2, or 1.5× acuity). For the amblyopic children, three were tested with sizes 3× acuity thresholds, seven with 2.5×, five with 2×, and five with 1.5×. For those with typical vision, one child was tested with 2.5×, six with 2×, and thirteen with 1.5×.

#### Adults

Adults completed the same three tasks as the children with the same stimuli (minus reward animations), with the addition of a Gaussian fixation point near the bottom of the monitor. Stimuli were presented monocularly to the dominant eye and viewed peripherally at four eccentricities: 2.5°, 5°, 10°, and 15° in the upper visual field.

On each trial of the acuity and crowding-extent tasks, the fixation point first appeared for 500ms. This was followed by the target, either in isolation (acuity task) or surrounded by the ghost flankers (crowding-extent task) for 500ms. A circular 1/f noise mask with a diameter of 1/3 the target eccentricity was then presented for 250ms. A different mask was presented on each trial. After the presentation of the mask, observers had unlimited time to make a 4AFC response on the target orientation. A 500ms inter-trial interval followed, with the fixation dot on screen. Each staircase consisted of 45 trials, with observers completing two staircases per eccentricity. For each observer, acuity and crowding-extent values were taken from the average gap-size threshold across the two staircases at each eccentricity.

The orientation-matching task was largely identical to the children’s version (Figure 1E). Stimuli were presented with a gap size of 3× the acuity threshold. With a target-flanker separation of 1.1× the target diameter, this gave absolute target-flanker separations that increased with eccentricity, averaging 0.64° at 2.5° eccentricity (range= 0.57-0.72°), 1.01° at 5° (range= 0.93-1.12°), 1.91° at 10° (range= 1.39-2.28°), and 3.13° at 15° (range= 2.39-4.41°). These values fell within the crowding extent measured for each observer. Average values were also close to being a constant proportion of the target eccentricity, corresponding to 0.26×, 0.20×, 0.19×, and 0.21× the target eccentricity, respectively. Besides the slight elevation at 2.5° eccentricity, the strength of crowding should therefore have been similar at all eccentricities.

The trial presentation sequence was similar to the acuity and crowding tasks, with a fixation dot appearing for 500ms, followed by the target for 500ms. The target was either presented in isolation or surrounded by flankers of a ±30° or ±90° orientation difference. A 1/f noise mask was then presented for 250ms, at which point a reference stimulus identical to the target appeared at fixation at a random orientation. The size of the reference matched the target. Observers had unlimited time to adjust the reference stimulus to match the orientation of the previously presented target. Adults completed 5 blocks of 100 trials per eccentricity, resulting in a total of 2000 trials per observer. In each block, 20 trials were included for each of the 5 flanker conditions. Blocks for each eccentricity were interleaved to counter any practice effects. Observers received auditory feedback in the form of a beep when their estimate of the target orientation was offset by more than ±35°. All other parameters were identical to the children’s version of the tasks.

## Results

### Acuity and Crowding Extent

The acuity and crowding-extent tasks each gave a measure of the gap size of the Vac-Man target required for performance to reach a particular point (62.5% for acuity; 80% for crowding). For acuity, the gap size was the value of interest. For crowding, the spatial extent value was calculated as the radius from the centre of the target to the centre of one flanker. This centre-to-centre separation was equal to the target diameter (which was five times the gap size) multiplied by 1.1 (the relative separation between elements). Here we consider the adult results first, followed by the children.

#### Adult Periphery

Gap-size (acuity) thresholds for the four eccentricities can be seen in Figure 2A. Thresholds increased with eccentricity, with 2.32±0.07 arcmins (mean±SEM) at 2.5° eccentricity, 3.68±0.07 arcmins at 5°, 6.95±0.34 arcmins at 10°, and 11.36±0.72 arcmins at 15°. A one-way ANOVA revealed a significant effect of eccentricity, (F[1.47, 13.20] = 110.34, *P*< .0001, Greenhouse-Geisser corrected), demonstrating the well-known reduction of acuity in the periphery.

**Figure 2.**
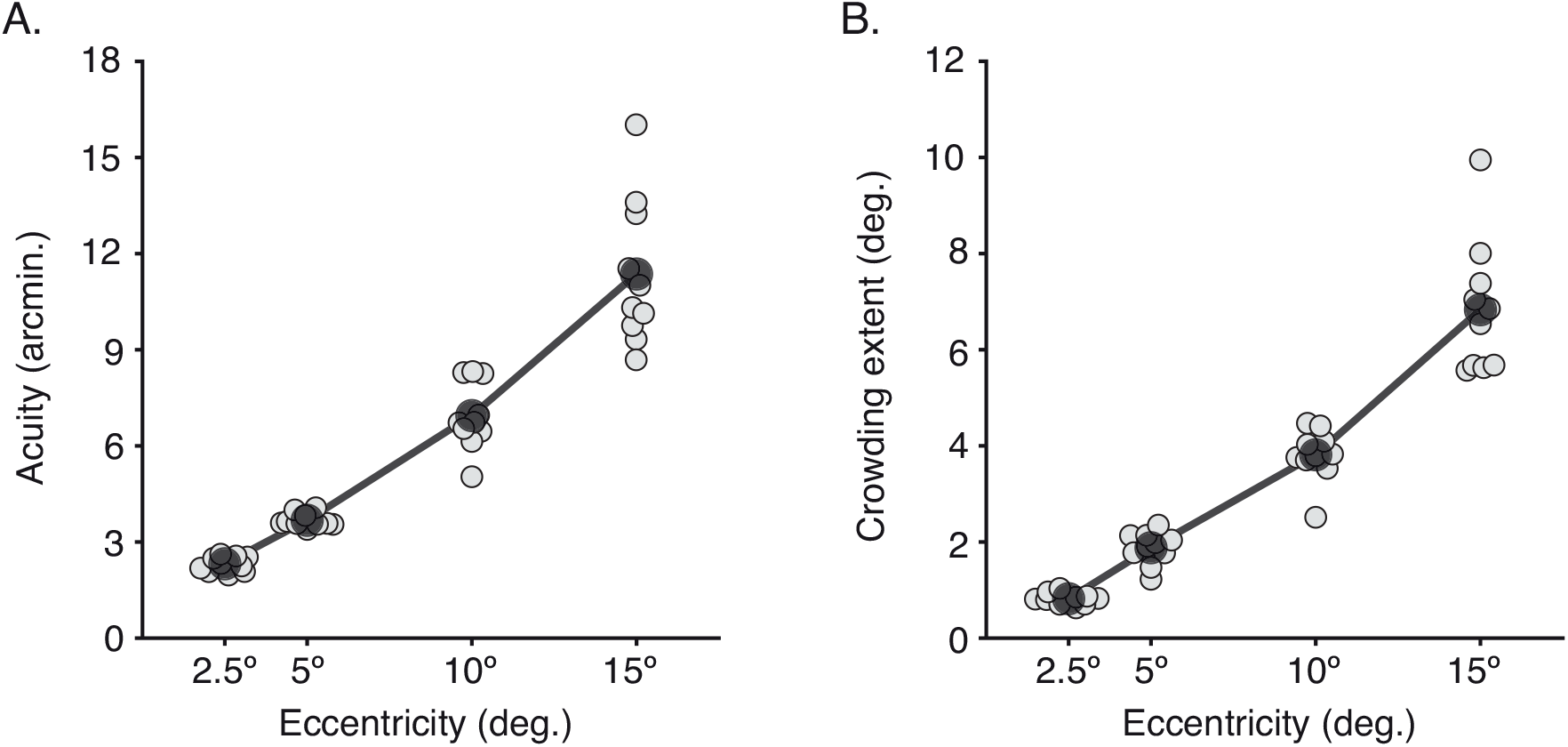
Acuity and crowding in the adult periphery. **A**. Acuity measured as gap-size thresholds (in minutes of arc) for adult observers at four eccentricities in peripheral vision. Light grey dots plot values for each observer (n=10), shifted along the x-axis where required for visibility, with the mean for each eccentricity overlaid as a transparent black dot. **B**. The spatial extent of crowding, measured as the centre-to-centre separation between the target and flankers in degrees of visual angle, plotted as in panel A.

The spatial extent of crowding at each eccentricity is presented in Figure 2B. Crowding also increased with eccentricity, though with a vast difference in scale from acuity, with 0.82±0.04° (mean±SEM) at 2.5°, 1.88±0.11° at 5°, 3.81±0.17° at 10°, and 6.83±0.44° at 15° eccentricity. A one-way ANOVA accordingly revealed a significant effect of eccentricity (F[1.23, 11.07] = 146.20, *P*< .0001, Greenhouse-Geisser corrected).

#### Typically Developing and Amblyopic Fovea

Acuity values for children with typical vision and amblyopia are plotted in Figure 3A. Gap-size thresholds for the left and right eyes of children with typical vision averaged 1.07±0.07 and 0.94±0.06 arcmins respectively (close to a Snellen acuity of 6/6). There was no significant difference between these values (paired samples t-test: t[19]=2.37, *P*= .5), indicating no interocular differences in acuity. Reduced acuity levels were evident in the amblyopic eye of the amblyopic group, with an average of 4.03±0.67 arcmins compared to 1.14±0.14 arcmin for the unaffected fellow fixating eye (equivalent to Snellen acuities of 6/24 and 6/6). This interocular difference was significant (paired samples t-test: t[19] =4.13, *P*< .001), consistent with the characteristic acuity deficit in amblyopia. Acuity in the fellow eye did not differ from the acuity of the children with typical vision (unpaired t-test between the fellow fixating eye and the mean of both eyes in children with typical vision: t[38]= -0.88, *P=* .38).

**Figure 3.**
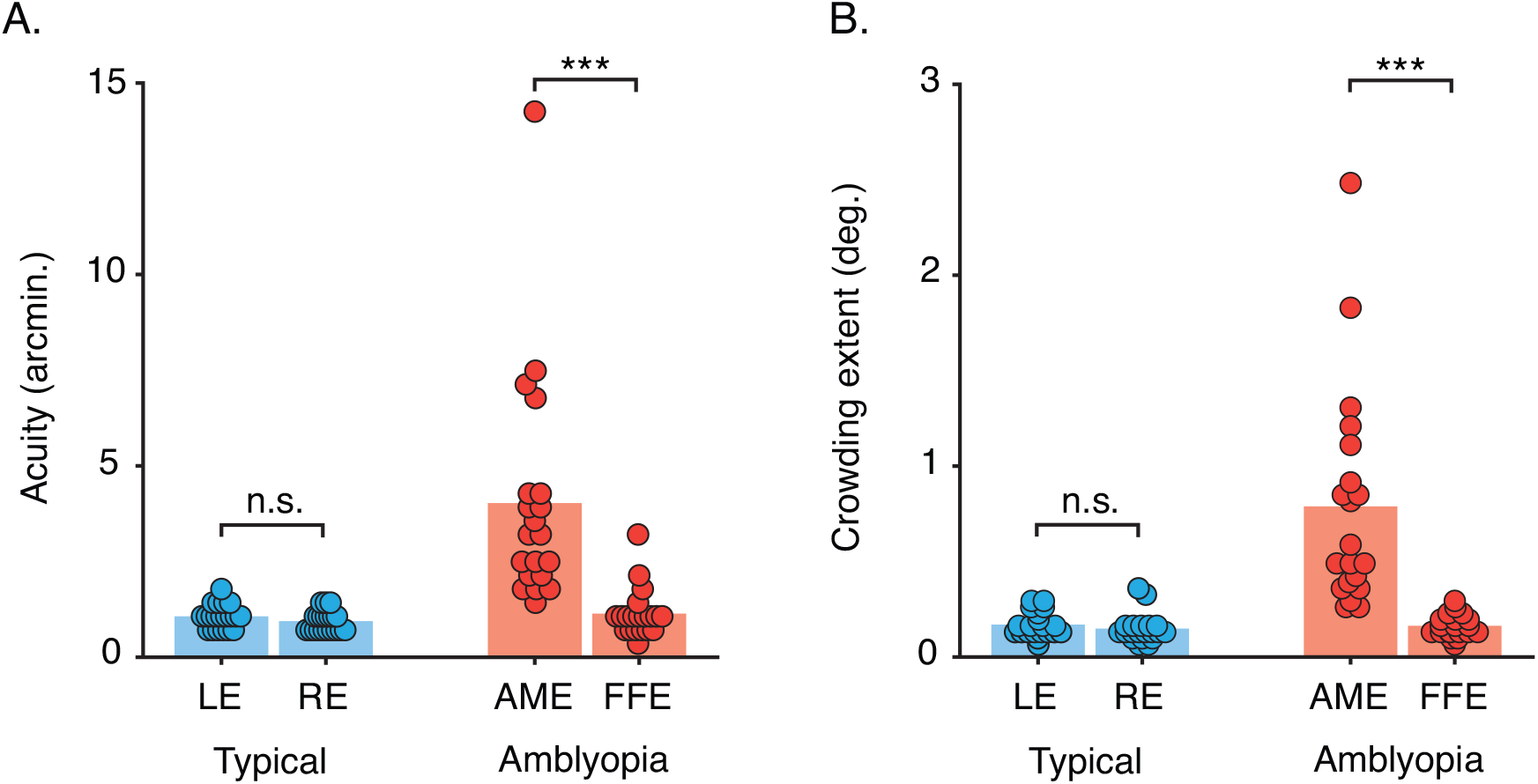
Acuity and crowding in the typically developing and amblyopic fovea. **A**. Acuity measured as gap-size thresholds (in minutes of arc) for children with typical vision and amblyopia (n=20 each). Dots indicate values for each eye of individuals (shifted on the x-axis for visibility); bars indicate the mean. LE = left eye, RE= right eye, AME= amblyopic eye, FFE = fellow fixating eye, n.s. = no significant difference, ***P<.001. **B**. The extent of crowding (in degrees of visual angle) measured as the centre-to-centre separation between the target and flankers, plotted as in panel A.

Values for the spatial extent of crowding for children are in Figure 3B. For children with typical vision, the extent of crowding averaged 0.17±0.01° for the left eye and 0.15±0.02° for the right eye, with no significant interocular difference (paired samples t-test: t[19]=1.58, *P*= .13). For the amblyopic group, the extent of crowding was greater in the amblyopic eye, averaging 0.79±0.13° compared to 0.16±0.01° for the fellow fixating eye (paired samples t-test: t[19]=4.85, *P*< .001). There was no difference in the extent of crowding between the fellow fixating eye of the children with amblyopia and the mean of both eyes in children with typical vision (unpaired t-test: t[38]=-0.21, *P*= .83).

### Orientation-Matching Task

Responses in the matching task were recorded as the perceived orientation of the target on each trial, which were subtracted from the veridical target value to give error values. Frequency histograms were constructed to tally the errors from ±180° in 10° bins, separately for each flanker condition. For children, this gave five distributions of response errors per observer. For adults, responses were combined across the five repeat blocks to give five distributions for each eccentricity. Both children and adults produced error distributions that were mirror-symmetric in conditions with target-flanker differences of opposite sign (−30°and 30°, -90° and 90°). As a result, the sign of the response errors was reversed in conditions with negative differences in order to sum the distributions. For each group, this gave three response-error distributions per observer (and per eccentricity for adults): unflanked, ±30° target-flanker difference, and ±90° target-flanker difference.

#### Adult Periphery

Figure 4 plots response-error histograms for each eccentricity in the adult periphery. For unflanked targets (left column), the distribution of response errors was unimodal across all eccentricities (panels A-D), with a peak at 0° and a narrow width. As such, when the target was presented in isolation, observers reported its orientation with good accuracy and precision, with increasing eccentricity having no effect on these estimates.

**Figure 4.**
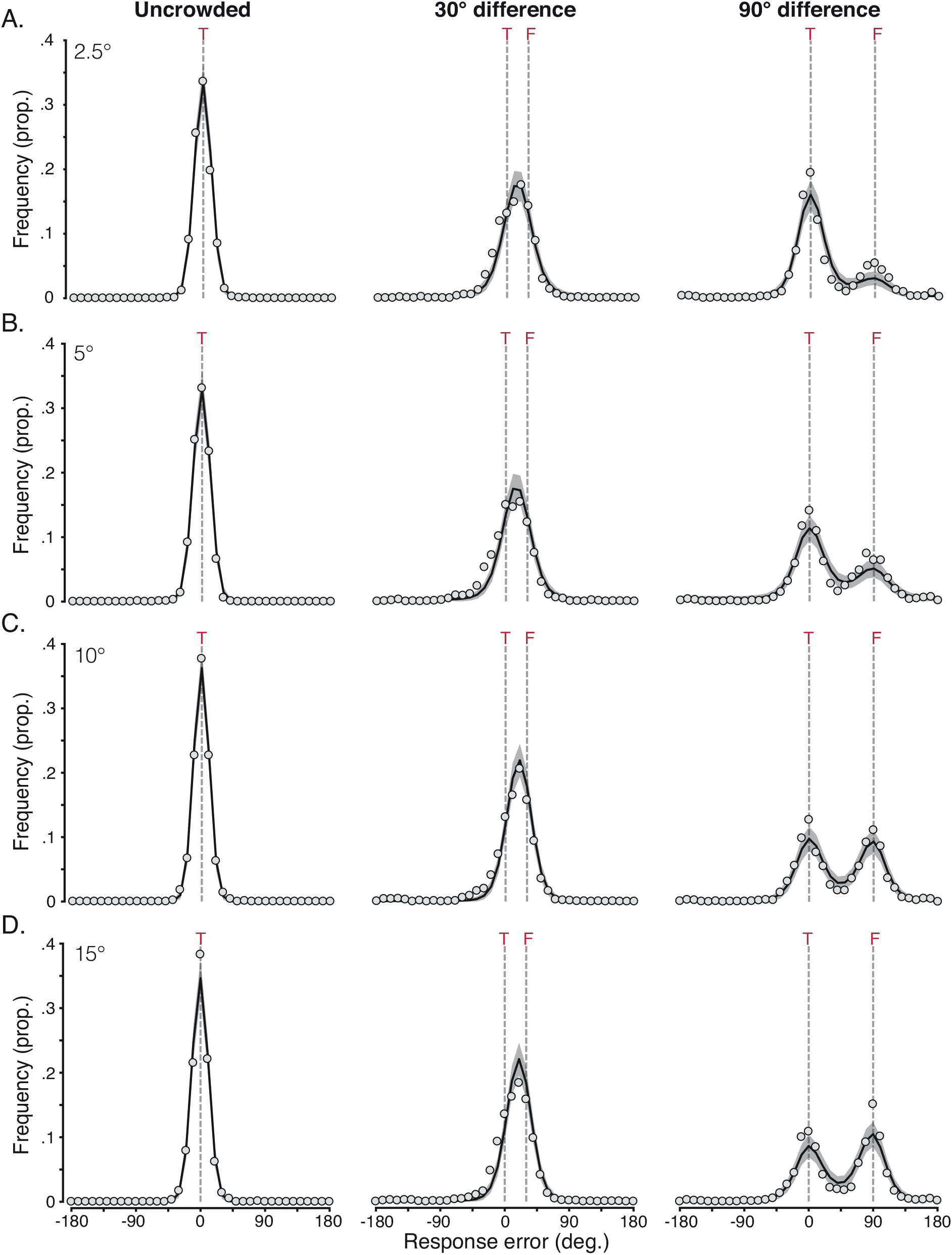
Distributions of mean response error from the orientation-matching task for the adult periphery. **A**. Response error distributions at 2.5° eccentricity, with mean values presented as light-grey dots. The black solid line plots the mean distribution of the population pooling model, with grey shaded areas plotting the 95% range of simulated distributions for 1000 model iterations. Dashed grey lines indicate the target location (‘T’), and for the conditions in which flankers were present, the flanker location (‘F’). **B-D**. Response error distributions at 5°-15° eccentricity, plotted as in panel A.

With flankers that differed by 30° (middle column), response-error distributions were also unimodal at all four eccentricities, though distributions shifted towards the flankers, with a peak at orientations between the target and flanker values (shown as dashed lines). There was also an increase in the spread of response errors relative to the unflanked condition. In other words, crowding had a disruptive effect on both accuracy and response precision.

When flankers differed by 90° from the target, response-error distributions became bimodal (Figure 4, right column). The first peak was concentrated at 0°, indicating responses near to the veridical target value, with the second at 90°, near to the flanker orientation. The location of these peaks did not change with eccentricity, though the height of the peaks did. At lower eccentricities, the frequency of responses near the target was greater (i.e. the peak centred on 0° was highest), whereas at larger eccentricities responses near the flanker orientation became more frequent. In other words, observers were increasingly likely to report orientations near to the flankers as eccentricity increased.

These response-error distributions allow us to draw a number of conclusions regarding the perceptual effects of crowding in the adult periphery. Observers were both accurate and precise when reporting the orientation of unflanked targets. With ±30° target-flanker differences, crowding primarily led observers to indicate intermediate orientations between the target and flankers. These responses can be classified as assimilation errors. Crowding with ±90° target-flanker differences led to a mixture of responses near to either the target or flanker orientations. The latter can be classified as substitution errors, which increased in frequency with eccentricity.

#### Typically Developing and Amblyopic Fovea

Figure 5 shows histograms of the response errors for children. When the target was unflanked, children with typical vision (Figure 5A, left) gave a unimodal distribution of response errors centred near 0°, with a slightly broader width than that found with adults in the periphery (Figure 4). The group with amblyopia (Figure 5B, left) showed a similar pattern of errors when stimuli were presented in the fovea of their amblyopic eye, with the peak centred on 0° and a similar bandwidth to children with typical vision. In other words, both groups could accurately report the orientation of the isolated target element with good precision.

**Figure 5.**
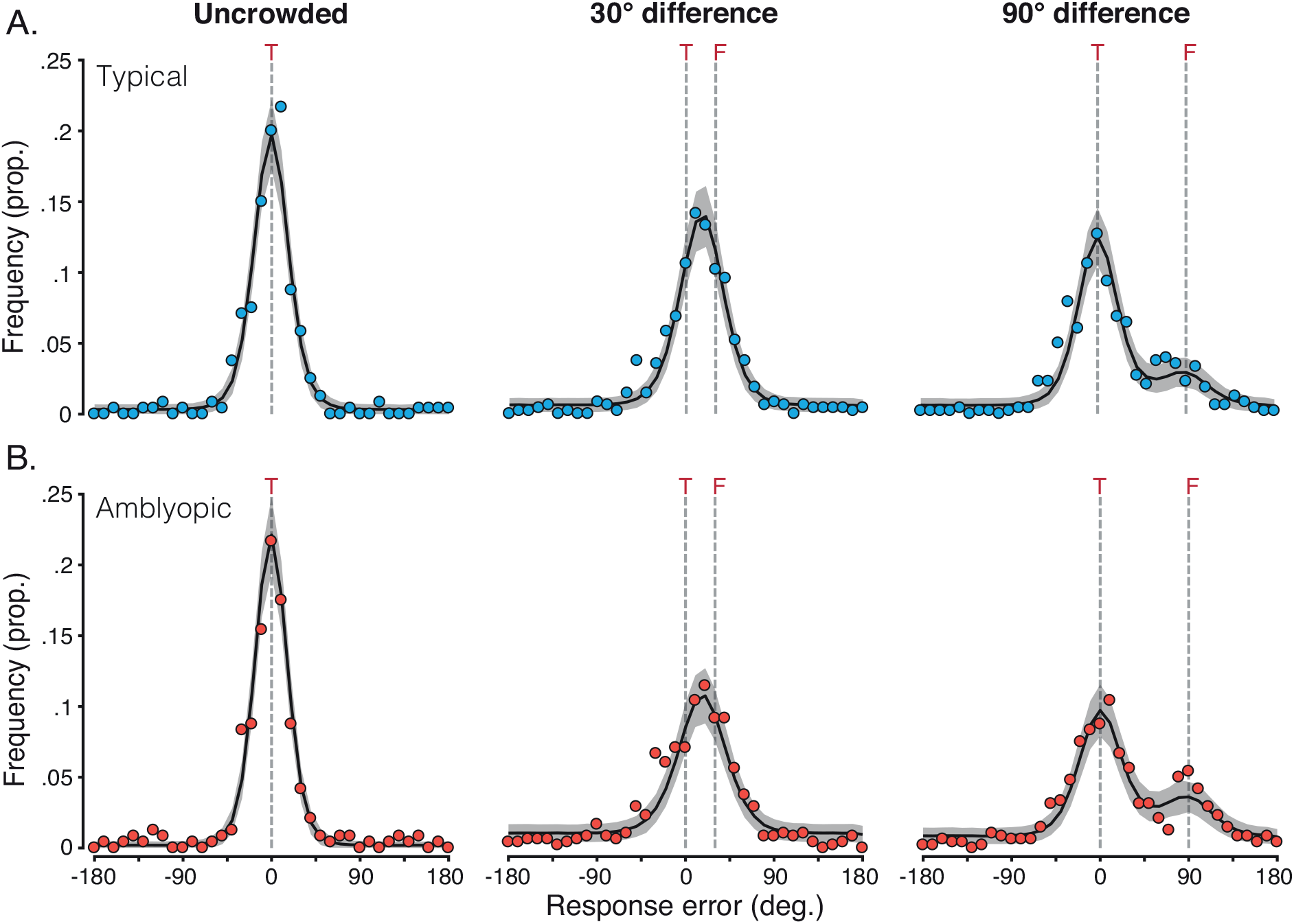
Mean response-error distributions for children with typical vision and amblyopia in the orientation-matching task. **A**. Response-error distributions for children with typical vision, with mean values shown as dots. The black solid line indicates the mean distribution of the population pooling model, with grey shaded areas plotting the range of simulated distributions for 1000 model iterations. Dashed grey lines indicate the target orientation (T), and when present, the flankers (F). **B**. Response-error distributions for children with amblyopia, plotted as in panel A. Note that all data here is for the amblyopic eye.

When flankers differed from the target by 30°, both groups of children showed unimodal response-error distributions with a peak shifted to fall between the veridical target and flanker orientations. Both groups showed an increase in response-error variability compared to the unflanked condition, with greater variability in the amblyopic group. This shift in the peak of response errors, combined with reduced response precision relative to unflanked performance, is similar to that observed in the adult periphery.

When flankers differed by 90°, response-error distributions for both groups of children became bimodal. In each case, responses were most frequently near to the target orientation, with a secondary peak in responses close to 90°, indicative of flanker reports. These flanker responses were more frequent in children with amblyopia. There was also an increase in the variability of responses relative to unflanked performance, but to a lesser extent than when the flankers differed by 30°. These response-error distributions were highly similar to those of adults in the near periphery (Figure 4 A&B).

Taken together, children performed well when required to judge the orientation of unflanked targets, giving responses that were both accurate and precise. With ±30° target-flanker differences, errors were primarily reports of intermediate orientations between the target and flankers. As with adults in this condition, these can be classified as assimilation errors. With ±90° target-flanker differences, children primarily reported values near to the target orientation, with a secondary rate of responses to the flanker orientation. The latter can be classified as substitution errors, which arose at a similar rate to the errors made by adults in the near periphery. Therefore, on a group level, children with typical vision and amblyopia made the same systematic errors with stimuli viewed foveally as did adults with stimuli viewed peripherally.

### Modelling

Given the common pattern of crowded errors made by adults and children with both typical vision and amblyopia, we next sought to examine the basis of these systematic perceptual outcomes with a computational model. Our approach was inspired by the models of van den Berg, Roerdink, and Cornelissen (2010) and Harrison and Bex (2015), and similar to recent models of crowding for colour and motion (Greenwood & Parsons, 2020). The model had three stages, summarised in Figure 6.

**Figure 6.**
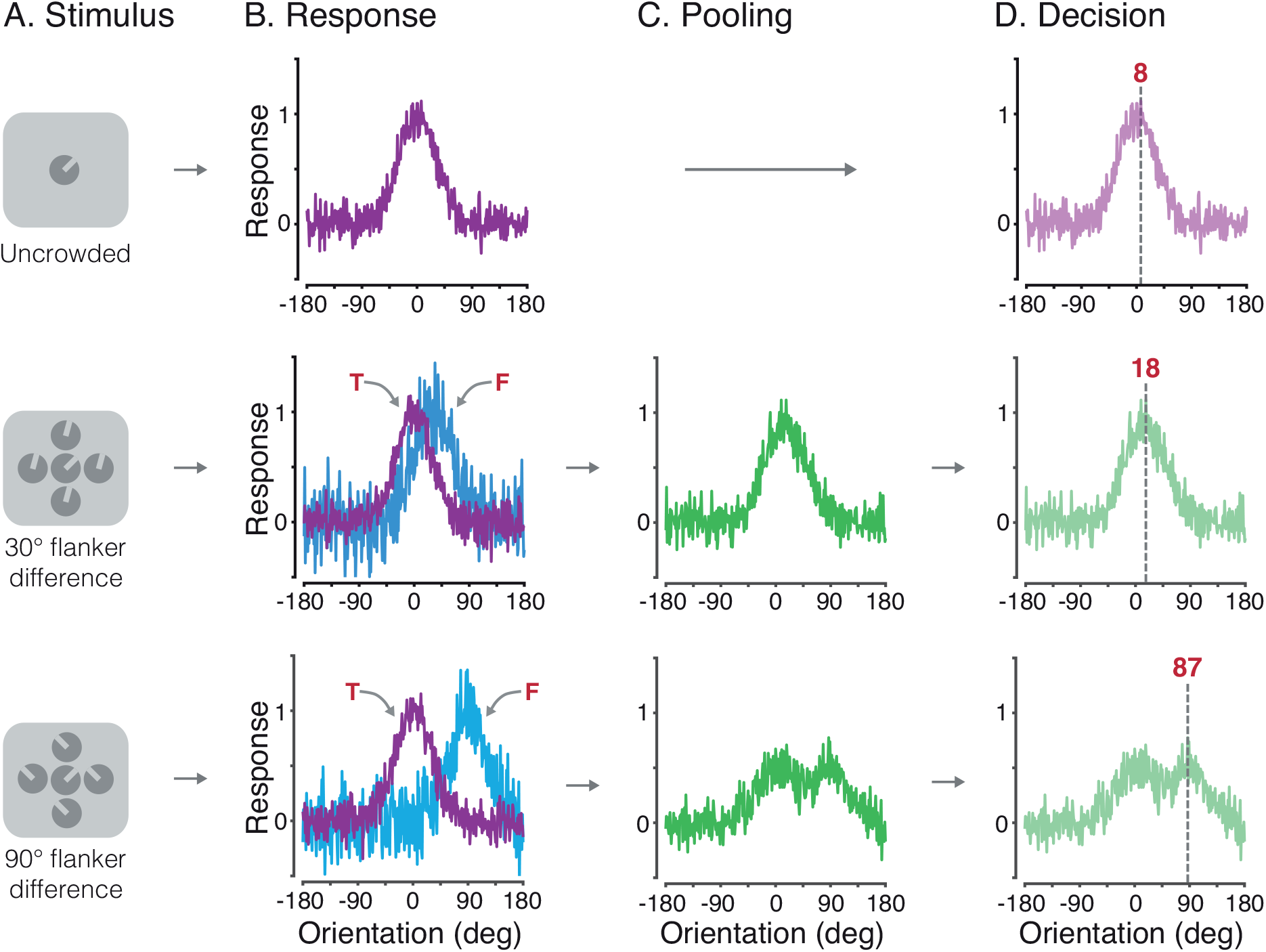
Illustration of the 3 stages of the weighted population pooling model. **A**. Example stimulus input for each of the 3 flanker conditions (unflanked, and flankers with a 30° or 90° difference from the target). **B**. Early population response to the target (upper panel), and the target and flankers (middle and bottom panels). Arrows indicate the response to the target orientation (‘T’; purple distributions) and the response to the orientation of the flankers (‘F’; blue distributions). **C**. The pooling stage, modelled as the weighted combination of population responses to the target and flankers. **D**. The decision stage, where the perceptual outcome is read out as the peak of the combined population response. The grey dashed line indicates this peak, with the value shown numerically.

Figure 6 A shows example stimuli for each condition. For ease of modelling, response errors were directly simulated using the orientation difference from the target as inputs (rather than absolute orientations). As in the results presented above, 0° indicated no error. In the first stage of the model (Figure 6B), the response of a population of detectors selective for orientation was simulated, similar to neurons in primary visual cortex (Schiller, Finlay, & Volman, 1976). Each detector responded to a range of orientations, according to a Gaussian tuning function with a peak sensitivity centred on a particular orientation, as in equation 1:

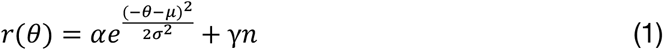

Here, *r(θ)* is the population response at a given orientation (θ), ranging from ±180°. The value α set the height of the detector sensitivity (set to 1), µ set the orientation producing the peak response, and σ gave the bandwidth, set to 30° to match the selectivity of neurons in cortical area V1 (De Valois, Yund, & Hepler, 1982). Gaussian noise *n* was added with a magnitude of γ (the first free parameter). This ‘early noise’ allowed us to fit sensitivity to unflanked orientations in particular, though the same noise parameter was used across all conditions.

Based on the principles of population coding (Pouget, Dayan, & Zemel, 2000), the resulting population-response distribution is a Gaussian function centred on the orientation of the Landolt-C stimulus, with a bandwidth equivalent to the underlying sensitivity bandwidth of the detectors. On trials when the input to the model was a target presented in isolation (unflanked), the resulting population responses were centred near 0°. Responses from an example unflanked trial are shown in Figure 6B (top panel). For illustration, in this example trial the early noise is set to 0.1.

In the 30° and 90° flanker difference conditions, population responses were generated for both the target *θ*_*t*_ and the flanker orientation *θ*_*f*_ on each trial. Flanker responses were generated using equation 1, though with a second free parameter for the noise term (‘late noise’), which allowed us to determine the degree of performance impairment induced by the flankers. The resulting population responses to the flanker orientation were centred either near 30° or 90°, respectively. Figure 6B shows population responses from an example trial for each of the two flanker conditions (middle and bottom rows), with the late noise parameter set to 0.2.

The second stage of the model simulated the effects of crowding as the pooling of population responses to the target and flanker elements. A weighted sum of the population response to the target and flankers was taken that allowed modulation of the precise combination of these responses. The weighted combination of responses to the target and flankers was given as:

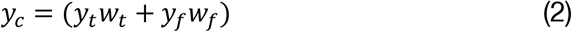

Here, *w*_*t*_ and *w*_*f*_ were the weights for the population responses to the target and flankers, respectively. The flanker weight ranged from 0-1, with the weight of the target being one minus the flanker weight. As the data from all groups indicated that response-error distributions in the 30° flanker difference condition differed from those in the 90° flanker difference condition, the flanker weight was independent in these conditions. This gave two additional free parameters: the flanker weight for the 30° condition 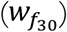 and that for the 90° condition 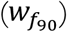, to give four free parameters in total.

Figure 6C shows the second stage for each flanker condition on an example trial. With a 30° target-flanker difference and a flanker weight of 0.5, both target and flanker distributions contribute equally to the pooled response. Their breadth means that the combined population response distribution is broadly unimodal, with a peak that shifts away from 0° towards the 30° flanker orientation. With a 90° target-flanker difference and the same 0.5 flanker weight, the combined population response distribution becomes bimodal, with peaks near the target and flanker orientations.

In the third and final stage of the model, a ‘decision’ on the perceived target orientation was made by extracting the maximum response from the population on each trial. For unflanked targets (Figure 6D), the population response to the target carried through to the final stage, and the peak of the response is near the target at 8°. For the 30° and 90° flanker difference conditions, the peak of the combined target and flanker population responses was taken. In the 30° flanker condition, this example trial gave a shift in the peak towards the flankers (at 18°) – an assimilation error. A decrease in the flanker weight here would shift the peak back towards the target orientation (0°), and vice versa. In the 90° flanker condition, the combination of the bimodal distribution and noise results in a peak closer to the flankers (at 87°) – a flanker substitution error. A decrease in the flanker weight here would increase the likelihood of responses laying around the 0° target value, rather than around the 90° flanker.

Our model simulation included 1000 trials per flanker condition. For comparison between these simulated responses and the measured error distributions, the output of the model was binned in 10° increments. This binning did not alter the output of the model in a qualitative fashion. The best-fitting parameters for all groups were determined using a two-stage fitting procedure. The initial coarse fit involved a grid search through the parameter space in pre-defined steps. From this we derived the parameters that best fit the data in the grid using the least-squared error (LSE) between response-error distributions and the simulated distributions from 1000 trials, summed across all 3 conditions. In the second-stage fine fit, the best parameters from the coarse fit were used to seed the analysis, with the best-fitting parameters determined by minimising the LSE using *fminsearch* in Matlab, again taking the summed difference between data and simulated distributions from 1000 trials. We then ran 1000 iterations of the model with the best-fitting parameters for each dataset.

We also considered the response errors that would arise if crowding does not have a systematic effect on target appearance, but rather distorts or adds noise to the target orientation. To do so, we tested an additional model that did not contain the second pooling stage, but rather simply added noise to the population response when flankers were present. Although distortions may be locally systematic (Pugh, 1958; Sireteanu, Lagreze, & Constantinescu, 1993), in the present task this would cause errors when the target and/or flanker gaps were rotated to appear in some visual field locations and not others. The variety of orientations and target-flanker combinations in our task would then produce a random distribution of errors across the experiment as a whole, particularly when errors were aligned to the target orientation rather than their absolute orientation (since distortions should depend on the retinotopic location of the gap in each element), and even more so once errors were pooled at the group level given the idiosyncrasy of these distortions. Errors due to attentional lapses or developmental issues with short-term memory (Witton, Talcott, & Henning, 2017; Manning et al., 2018) should similarly manifest as random errors.

The noise model had three stages, the first of which was identical to the population pooling model, using equation 1 above to produce a population response to the target. Early noise was again the first free parameter. Population responses to the flankers were not simulated. Rather, the second stage differed in that the population response to the target was simply subjected to an additional noise parameter:

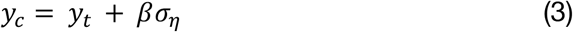

Here, y_t_ is the population response to the target, σ_η_ the added random noise and *β* was the magnitude of this late noise (the second and final free parameter in the noise model). The third stage of the model was then identical to the final stage of the pooling model, with the maximum response taken as the ‘decision’ of the model on each trial. As with the population pooling model, we ran 1000 trials of this 3-stage model for each condition, using the procedure described above. MATLAB code for both models is available online^i^.

### Model Simulations of Group Data

#### Adults

Figure 4 shows the result of 1000 iterations of the best-fitting population pooling model for the adult mean data at each eccentricity. For unflanked targets, the model almost perfectly captures the response errors, with early noise values that were similar across all eccentricities (see Table A4, and plotted in Figure 7 as white triangles). When the target was crowded, response distributions became noisier and more broadly distributed, with a lower peak response. The model captured this through the addition of late noise to the flanker population response, which added further disruption to the pooling stage. Values for the late noise parameter showed a slight decrease with eccentricity, from values of 1.22 at 2.5° to 0.73 at 15° eccentricity.

**Figure 7.**
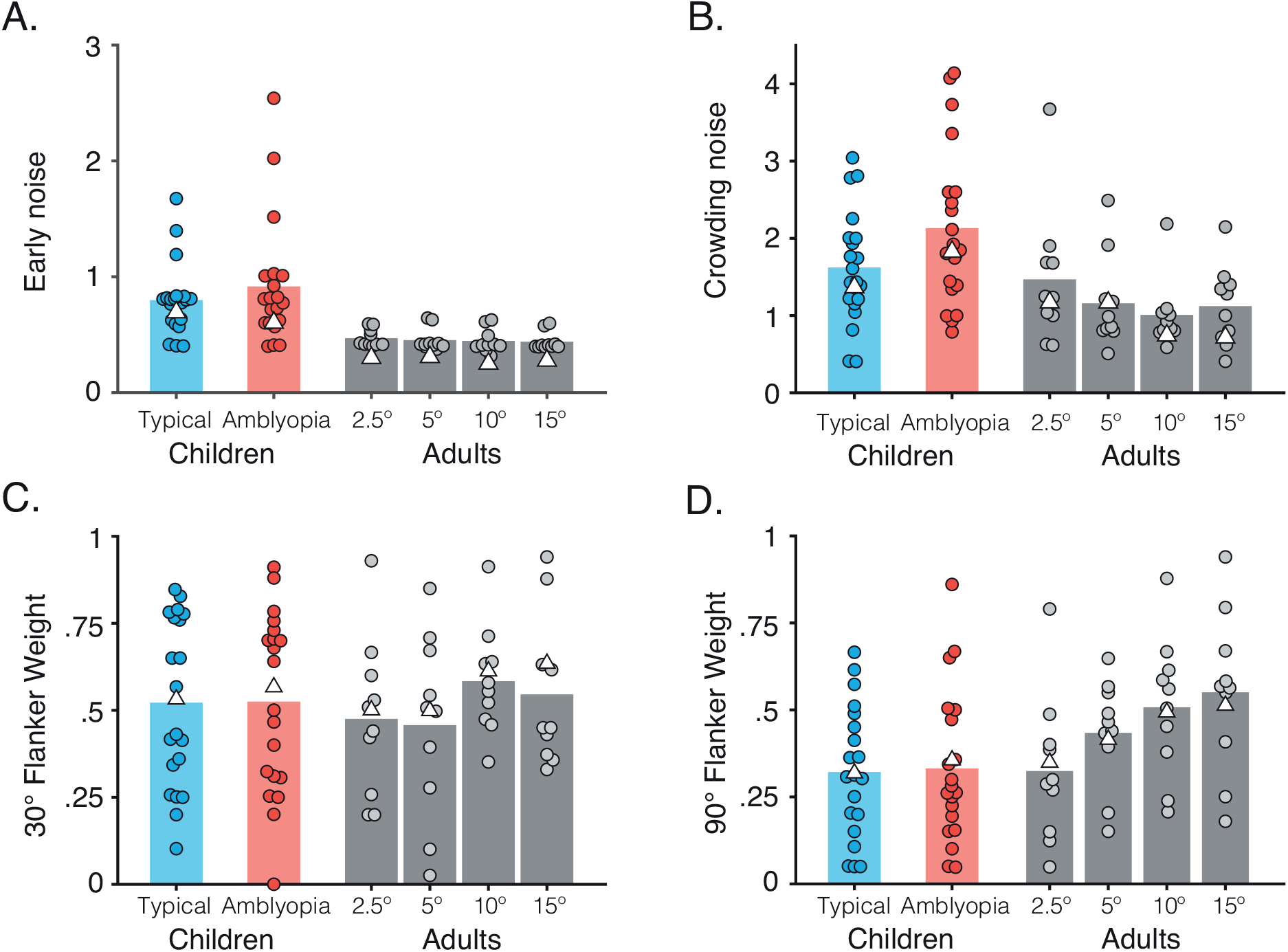
Best-fitting model parameters in the adult periphery and for children with typical vision and amblyopia. **A**. Best-fitting values for the early noise parameter. Dots indicate individual observers (shifted on the x-axis for clarity), and bars the mean of the individual observers. White triangles show the parameters for fits to the averaged group data. **B**. Best-fitting values for the late noise parameter, plotted with conventions as in A. **C-D**. Best-fitting values for the flanker weights when the flankers differed by 30° from the target (C),and when flankers differed by 90° (D).

The systematic nature of the response errors induced by flankers were driven by the flanker weight parameters. With ±30° target-flanker differences, the model clearly replicates the response-error distribution in Figure 4 (middle column) where the majority of the errors were between the target and flanker orientations. These flanker weights increased with eccentricity from 0.52 at 2.5° eccentricity to 0.64 at 15°. With ±90° target-flanker differences, there was a marked effect of eccentricity, with the proportion of flanker responses increasing with eccentricity. This was captured well by the model, which followed this pattern of increasing flanker responses with an increase in the flanker weights from 0.34 at 2.5° to 0.52 at 15° eccentricity. On the whole, the model follows the profile of response errors in all conditions.

In contrast, the noise model generally performed poorly, as shown in Figure A1 of Appendix B. Although the model was able to capture errors around the target in the unflanked condition, it failed to produce the shift towards the flankers in the 30° condition and the bimodal pattern of errors in the 90° condition. Clearly the addition of noise alone, ignoring the identity of the flankers, is insufficient to account for these errors. To determine which of the two models best fit the response distributions, we computed Akaike Information Criterion (AIC) values (Akaike, 1974) which take the least squared error (LSE), and correct for the number of parameters included in each model (since the pooling model had four free parameters while the noise model had two). Despite this correction, the superior performance of the population pooling model is clear from the AIC values, shown in Figure 8A (and Tables A3 & A4), where lower AIC values indicate better model fits to the data. In adults, the pooling model outperformed the noise model at all eccentricities. As such, the group response errors could not be accounted for by a process that merely adds random noise to the target orientation.

**Figure 8.**
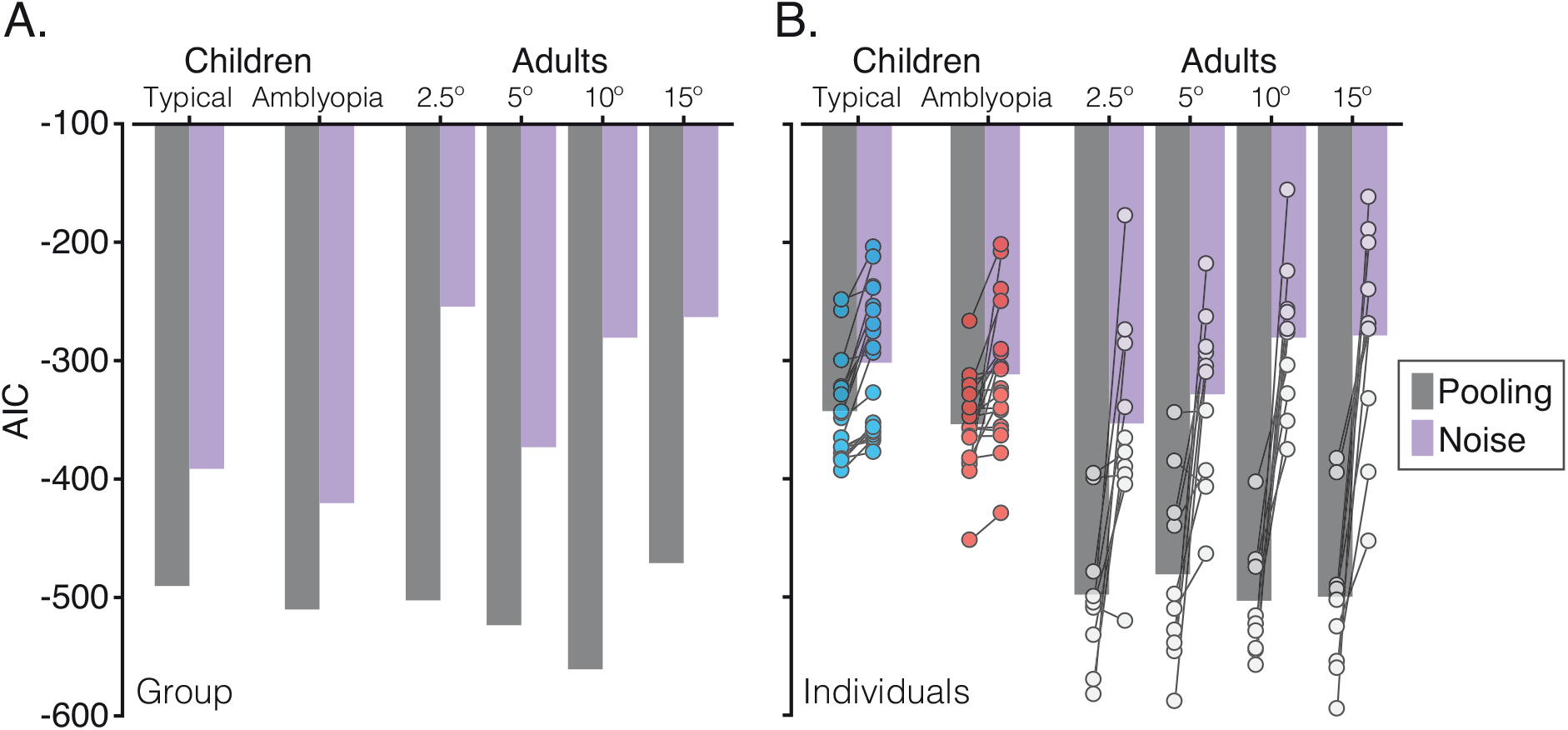
Akaike Information Criterion (AIC) values to assess goodness-of-fit for the pooling and noise models. **A**. AIC values for the fits to the averaged group data, plotted separately for the pooling (dark grey) and noise (light grey) models. Note that more negative values indicate better fits to the data. **B**. AIC values for model fits to individual data. Bars here represent the mean of the fits to individual data for the pooling and noise models, with individual values superimposed as circles. Individuals are joined with bars to show the direction of effect.

To reproduce the systematic shifts in these error distributions, the pooling model used two additional free parameters for flanker weights in the ±30° and ±90° flanker conditions. In Appendix C we compare this with the performance of a three-parameter version of the pooling model, where the same weight was applied in both flanker conditions. Although this three-parameter model performed better than the noise model (see Table A5), it was less effective at capturing the systematic pattern of crowded errors than the four-parameter pooling model (Figure A2). It is thus important that these models incorporate variations in the strength of crowding with target-flanker differences in orientation.

#### Children

Figure 5 shows the simulated distributions for the group data of children with typical vision (Figure 5A) and amblyopia (Figure 5B), each computed as the mean of 1000 model iterations. In the unflanked condition, the model captures the Gaussian distribution of response errors well for both groups, with early noise values higher for both groups than those of adults (Table A4). When the target was crowded, the combined population response again became noisier and more broadly distributed, with a lower peak response. This was again captured by the model with the addition of late noise, values of which were again higher than those used for adults.

To reproduce the shift in response error distributions for the crowding condition with ±30° target-flanker differences, flanker weights in the model were similar to those of the two closest eccentricities in adults. With ±90° target-flanker differences, the model successfully captured both peaks of the bimodal distribution of response errors in each of the two groups, again using weights similar to adult values in the parafovea. Overall, the model successfully captured the pattern of systematic errors observed for both groups of children. The noise model again provided a substantially poorer fit to the data (see Appendix B and Figure A1), demonstrating that the errors of children with typical and amblyopic vision cannot be accounted for by a process that merely adds random noise to the target orientation. The four-parameter pooling model was similarly better able to characterise response errors than the three-parameter pooling model (Appendix C), again demonstrating the need for variations in flanker weights with target-flanker similarity.

### Model Simulations of Individual Data

Having demonstrated that the population pooling model can reproduce the observed response-error distributions using group data, we next consider how well the model can account for individual data. The model was fit using the same procedure as above, this time to data from individual observers. Because children had so few trials per condition (24 in each crowded condition), smoothing was applied to the response-error histograms using a three-point boxcar average prior to model fitting. Figure 7 shows the best-fitting values for all free parameters of the model for each adult at the four eccentricities tested, and for each child in the groups with typical vision and amblyopia.

For the early noise parameter (Figure 7A), values were again similar across the four eccentricities tested in adults, with mean values (shown as bars) between 0.44-0.47 and individual values from 0.32-0.64. These values were generally larger in children with typical vision (mean 0.80, range 0.40-1.68) and amblyopia (mean 0.92, range 0.40-2.54). For the late noise parameter (Figure 7B), adult values again showed a slight decrease with eccentricity on average, with means (and ranges) of 1.47 (0.61-3.67) at 2.5°, 1.16 (0.51-2.49) at 5°, 1.01 (0.59-2.19) at 10°, and 1.12 (0.41-2.15) at 15°. Children in both groups again tended to require higher late noise values, with a mean of 1.62 (range 0.4-3.04) for those with typical vision and 2.13 (range 0.79-4.14) for those with amblyopia.

When flankers differed by ±30° from the target (Figure 7C), flanker weights again tended to increase with eccentricity for adults, with means (ranges) of 0.48 (0.20-0.93), 0.46 (0.03-0.85), 0.58 (0.35-0.91) and 0.55 (0.33-0.94) from 2.5-15° eccentricity, though substantial individual differences are clearly apparent in the ranges. For children with typical vision and amblyopia, best-fitting flanker weights gave mean (range) values of 0.52 (0.10-0.85) and 0.53 (0.00-0.91), respectively, again corresponding to values in the adult periphery that were between those of the near periphery and the farthest eccentricities. When flankers differed by ±90° from the target (Figure 7D), flanker weights in the adult periphery showed a clear increase with eccentricity, with means of 0.32 (0.05-0.79) at 2.5°, 0.43 (0.15-0.65) at 5°, 0.51 (0.21-0.88) at 10°, and 0.55 (0.18-0.94) at 15°. The range was again broad, indicative of individual differences. In children, the range of flanker weight values was similarly broad, with means of 0.32 (0.05-0.67) and 0.33 (0.05-0.86) that were again most similar to the closer eccentricities in adults.

A number of conclusions can be drawn from the model fits to the individual data. First, children generally required larger early noise values than adults in order for the model to simulate their response error distributions. This indicates a broad difference in the general properties of foveal vision in children and the adult periphery. Children from both groups also required greater late noise values than adults, suggesting greater difficulties in clutter that may go beyond the stimulus, perhaps into decisional processes. Flanker weights in the children were however similar to values for adults in the parafovea, particularly for the 90° flanker condition, consistent with the commonalities observed in the pattern of response errors between adults and children.

In order to determine the success of these fits to individual data, the noise model was also fit to individual response-error distributions. Comparisons of these fits are shown in Figure A3 of Appendix D for adults and Figure A4 for children. Generally, as with the fits to the group data, the noise model failed to account for the systematic shift in the pattern of response errors towards the flanker orientations. However, for some individuals the noise model approached the success of the pooling model, and in others the noise model outperformed the pooling model.

AIC values for individual fits are shown in Figure 8B. In the adult periphery, the pooling model had lower AIC values than the noise model in 9 of 10 adults at 2.5° and 5°, and for all 10 adults at 10° and 15°. Accordingly, AIC values were significantly lower (indicating better fits) for the pooling model at all eccentricities: t[9]= -3.83, *P=* .004 at 2.5°; t[9]= -3.92, *P=* .004 at 5°; t[9]= -7.35, *P<* .001 at 10° and t[9]= -6.20, *P<* .001 at 15°. Those for whom the pooling model failed tended to have low error rates (as shown in Figure A3), making it hard to discriminate between the models.

For children, all 20 of those with typical vision showed lower AIC values for the pooling model, indicating better fits, while 18 of 20 amblyopic children were better fit by the pooling model. Here too the AIC values were significantly lower for the pooling model, with t[19]= -5.96, *P<* .001 for those with typical vision and t[19]= -3.67, *P=* .002 for those with amblyopia. Amongst the amblyopic children, those for whom the pooling model failed tended to have highly noisy responses (as shown in Figure A4), which again gave similar fits for the two models. Although this tendency was present in some of the children with typical vision, this was never to the extent that the noise model outperformed the pooling model.

On the whole, the pooling model outperforms the noise model in describing the response errors of all three groups. It is particularly striking that the pooling model is able to outperform the noise model with fits to individual data, given that these distributions relied on only 24 trials for children in each crowded condition. We conclude that the errors made by adults in peripheral vision cannot be accounted for by a process that merely adds random noise to the target orientation. The same is true for children with both typical and amblyopic vision. Rather, systematic changes in the target orientation are required to capture the systematic errors in these patterns of performance.

## Discussion

Our aim was to examine the perceptual effects of crowding in the fovea of children with typical vision and amblyopia, and to assess its similarity with crowding in the typical adult periphery. In all three groups, errors in the perceived orientation of a crowded Landolt-C target systematically followed the appearance of the flanker elements. With crowded targets, children made errors that predominantly matched either intermediate orientations between the target and flankers (assimilation) with 30° target-flanker orientation differences, or the orientation of the flankers (substitution) with 90° target-flanker differences (Figure 5). These errors matched those observed in adult peripheral vision (Figure 4), consistent with previous studies of adults (Parkes et al., 2001; Greenwood, Bex, & Dakin, 2009; Dakin et al., 2010; Ester, Zilber, & Serences, 2015; Harrison & Bex, 2015). The frequency of both assimilation and substitution errors increased with eccentricity in the typical periphery, with the errors made by children with typical vision and those with amblyopia most closely resembling the errors found at para-foveal eccentricities. In other words, children with typical vision and those with amblyopia make the same crowded errors as adults in the visual periphery. The commonality of these errors, in conjunction with other properties shared between these instances of crowding (Levi, Hariharan, & Klein, 2002a; Greenwood et al., 2012; Song, Levi, & Pelli, 2014) leads us to conclude that a common mechanism underlies crowding in these three instances.

We further demonstrate that a weighted population pooling model can reproduce this pattern of crowded errors in the typically developing and amblyopic fovea, as well as in the adult periphery. Similar to prior approaches (van den Berg, Roerdink, & Cornelissen, 2010; Harrison & Bex, 2015; Greenwood & Parsons, 2020), we simulated crowding via the weighted pooling of population responses to the target and flanker orientations. The weights determined the relative contribution of the target/flanker population responses to the pooled response distribution, from which decisions were drawn. With this approach, assimilation errors arose with 30° target-flanker differences because the combination of the target and flanker population responses gave a unimodal distribution with a peak response at intermediate orientations. Substitution errors arose with 90° target-flanker differences due to the bimodal distribution of the pooled response – although correct target responses were usually most common, noise in the bimodal pooled response gave a secondary rate of reports near the flanker orientation. Flanker population responses also added noise to the pooled population response, in line with previous models of peripheral crowding (Greenwood, Bex, & Dakin, 2009; Ester, Klee, & Awh, 2014; Ester, Zilber, & Serences, 2015; Greenwood & Parsons, 2020). However, noise alone was insufficient to account for the errors made by either group of children or in the adult periphery (Figure 8). Indeed, the success of the population pooling model was apparent not only in the fits to group data, but also with individual distributions of response errors. As a result, we suggest that pooling models provide a likely candidate for the common mechanism in these three instances of crowding.

In order to test the viability of the population pooling model, we have of course restricted our stimuli to simple target-flanker configurations (e.g. with identical flankers in a given trial). Nonetheless, given the success of pooling models in accounting for crowding with more complex stimuli in peripheral vision, including letters (Freeman, Chakravarthi, & Pelli, 2012) and faces (Kalpadakis-Smith, Goffaux, & Greenwood, 2018), it is likely that these approaches could similarly account for the crowding of complex stimuli in the developing and amblyopic fovea. The pooling process used herein is also consistent with higher-dimensional ‘texture’ pooling models, which can simulate crowding in natural scenes through the extraction of image statistics across large regions of the peripheral field (Balas, Nakano, & Rosenholtz, 2009; Freeman & Simoncelli, 2011; Keshvari & Rosenholtz, 2016; Rosenholtz, Yu, & Keshvari, 2019). Both assimilation and substitution errors have indeed been simulated with this approach in the adult periphery (Keshvari & Rosenholtz, 2016), making it likely that a similar process could work in the developing and amblyopic fovea, though it is unclear whether these models could also predict the increase in random responses found in children, as well as the individual differences observed.

The systematic effects of crowding that we observe are difficult to reconcile with higher-level models of crowding. For instance, attentional models (Strasburger, Harvey, & Rentschler, 1991; Strasburger, 2005) predict a predominance of substitution errors due to an inability to accurately focus spatial attention. The assimilation errors that we observe with 30° target-flanker differences are difficult to explain within this framework. Similarly, grouping theories propose that crowding is determined by Gestalt principles in a top-down fashion (Herzog, Sayim, Chicherov, & Manassi, 2015). These models fail to account for our data in two ways. First, they make no prediction regarding the systematic perceptual outcomes of crowding, focusing instead on the performance decrements induced by clutter. Second, their top-down operation is inconsistent with the monocular elevations in foveal crowding found in amblyopia – the same stimulus can cause crowding effects when presented to one eye, but not the other.

Consistent with prior reports of individual differences in the spatial extent of crowding (Toet & Levi, 1992; Petrov & Meleshkevich, 2011; Greenwood, Szinte, Sayim, & Cavanagh, 2017), our results also reveal substantial individual differences in the perceptual effects of crowding in both adults and children. For instance, with the 30° target-flanker difference, some typically developing children reported clearly intermediate orientations, whereas others reported orientations more similar to the flankers (see Figure A4 and Appendix D). The pooling model was nonetheless able to account for these individual differences through variations in the four free parameters. Although children showed substantial variations in these parameters relative to adults, some of this variation may be due to differences in age, given the developmental trajectory of crowding (Jeon et al., 2010), as well as variations in the extent of disruption from amblyopia.

Further evidence for a common mechanism for peripheral, developing, and amblyopic crowding comes from its featural selectivity. Both children and adults (in the near periphery) gave more responses near to the correct target orientation when target and flankers differed by ±90° than when they differed by ±30°. Our pooling model reproduced these differences with lower flanker weights for ±90° vs. ±30° flankers, which gave better fits to the data for all groups than a pooling model with a single weighting parameter for both conditions (Appendix C). This finding is consistent with the selectivity of peripheral crowding for target-flanker similarity in orientation, whereby errors decrease when elements are less similar (Andriessen & Bouma, 1976; Leat, Li, & Epp, 1999; Levi, Hariharan, & Klein, 2002b; Hariharan, Levi, & Klein, 2005). Here we show that target-flanker similarity also matters in developing and amblyopic crowding. This is inconsistent with prior reports that amblyopic crowding in adults disrupts target recognition regardless of target-flanker similarity (Hess, Dakin, Tewfik, & Brown, 2001; Hariharan, Levi, & Klein, 2005). Interestingly variations in featural selectivity are also evident in our results. At higher eccentricities in the adult periphery, the frequency of substitution errors increased with 90° target-flanker differences, suggesting that the selectivity for target-flanker similarity may decrease with eccentricity. This is also evident in the parameters of the pooling model (Figure 7), where flanker weights for the two conditions converged at higher eccentricities. Following suggestions that foveal crowding in amblyopia can be linked with an equivalent eccentricity in the typical periphery (Levi, Klein, & Aitsebaomo, 1985), we note that the developing and amblyopic fovea resemble para-foveal eccentricities more than the far periphery in terms of both the selectivity for target-flanker similarity and the systematicity of crowded errors.

Given the consistency of the response-error distributions in peripheral, amblyopic, and developing crowding, and the success of the population pooling model in reproducing them, we suggest that a common neural basis could underlie these effects. Because the weights in our pooling model are central to the crowding process, the question becomes: what determines the variations in these flanker weights? We suggest two possible factors. First, the increase in the extent of crowding with eccentricity (Bouma, 1970; Toet & Levi, 1992) has been attributed to increases in receptive field size (Parkes et al., 2001; Motter, 2009) and/or the overlap between receptive fields (Dayan & Solomon, 2010). Individual differences in crowding are indeed correlated with population receptive field (pRF) size in V2 (He, Wang, & Fang, 2019), with increased pRF sizes observed in the amblyopic fovea in areas V1-V3 compared to the typical fovea (Clavagnier, Dumoulin, & Hess, 2015). The role of receptive field size in developmental crowding is less clear, with neuroimaging studies suggesting that pRFs reach adult-like sizes by 6 years of age (Dekker et al., 2019), despite foveal crowding being elevated until as late as 12 years (Atkinson & Braddick, 1983; Atkinson et al., 1987; Jeon et al., 2010; Greenwood et al., 2012). It is possible however that the spatial selectivity of visual neurons may change in later childhood, given for instance the maturation of connections in primary visual cortex (Huttenlocher, de Courten, Garey, & Van der Loos, 1982; Huttenlocher & Dabholkar, 1997), and the later maturation of centre-surround receptive fields in extrastriate cortex (Zhang et al., 2005).

Alternatively, the rise in crowding with eccentricity has also been attributed to decreases in the cortical distance between target and flanker elements (Motter & Simoni, 2007; Pelli, 2008; Mareschal, Morgan, & Solomon, 2010) driven by cortical magnification (Daniel & Whitteridge, 1961; Van Essen, Newsome, & Maunsell, 1984; Sereno et al., 1995). Individual differences in cortical magnification may then explain variations in crowding, similar to proposals regarding variations in acuity (Duncan & Boynton, 2003) and perceived object size (Moutsiana et al., 2016). As well as spatial variations, the featural selectivity of crowding has also been argued to follow cortical distance – changes in orientation may shift target and flanker signals along orientation columns in visual cortex (Mareschal, Morgan, & Solomon, 2010), which could cause a reduction in weights in a similar manner to changes in target-flanker separation. Because amblyopia is known to produce a reduction in neurons responding to the amblyopic eye in areas V1 and V2 (Crawford & von Noorden, 1979; Bi et al., 2011; Shooner et al., 2015), the increased crowding with this condition could also derive from a reduction in cortical magnification. However, neuroimaging studies have thus far failed to observe differences in cortical magnification in either amblyopia (Clavagnier, Dumoulin, & Hess, 2015) or in the course of typical development (Dekker et al., 2019). Of course, our current data cannot distinguish between these effects of receptive field size/overlap and cortical distance. We suggest that one or both of these factors might drive the variations in weights that determine both the strength of crowding and its effect on the appearance of crowded objects.

An alternative explanation for these variations in crowding is that they may reflect differences in the positional uncertainty regarding the target element. Positional uncertainty could create source confusion (Wolford, 1975) either through the mislocalisation of features or whole letters (Strasburger, Rentschler, & Jüttner, 2011). A rise in positional uncertainty with eccentricity could in particular explain the rise in substitution-type errors found in the ±90° flanker condition. Indeed, both peripheral vision and amblyopic foveal vision are characterised by increased positional uncertainty (Hussain et al., 2015). Similarly, positional uncertainty may recede in the course of development, given the observation that Vernier acuity does not reach adult levels until the early teens (Carkeet, Levi, & Manny, 1997; Skoczenski & Norcia, 2002). As such, some proportion of the substitution errors found with 90° target-flanker differences may be attributable to position uncertainty in the adult periphery, as well as in children with amblyopia and typical vision. This is not necessarily inconsistent with pooling models of crowding, and indeed some have incorporated both processes (Freeman, Chakravarthi, & Pelli, 2012; Harrison & Bex, 2017). Our model also captures the increase in errors with ±90° flankers across eccentricity via an increase in the flanker weights for this condition, which could derive from positional uncertainty. This would however speak against the idea raised above that a common factor could drive all of the variations in crowding, as some have suggested (Agaoglu & Chung, 2016). Importantly however, position uncertainty alone cannot explain all of our results, given the predominance of assimilation errors observed with 30° target-flanker differences.

Despite the above-noted similarities between groups, differences are also evident. The model fits to data from both sets of children required higher levels of early noise in order to match performance, consistent with the broader elevations in low-level vision such as acuity and contrast sensitivity, both in amblyopia (McKee, Levi, & Movshon, 2003) and during development (Simons, 1983; Leat, Yadav, & Irving, 2009). This could also be driven by attention lapses or limitations of either short-term memory, motor coordination, or decision-making factors (Witton, Talcott, & Henning, 2017; Manning et al., 2018). A small subset of children with amblyopia were further noted to have made a disproportionate number of random reports (Figure A4), giving orientations that did not correspond to either the target or the flankers. For these cases, a model that simply adds noise to the target population response performed best (Figure 8). This increased response variability could be due to attentional lapses, or to perceptual distortions that characterise amblyopic vision (Pugh, 1958; Sireteanu, Lagreze, & Constantinescu, 1993; Barrett et al., 2003). More broadly however, these random factors cannot account for the systematic effects of crowding on target appearance, though it is possible that they acted as an additional source of errors, particularly for some children.

We have presented a population pooling model that can characterise the perceptual effects of crowding in three instances: the amblyopic fovea, the typically developing fovea, and the adult periphery. On the basis of this common mechanism, we can make predictions for a diverse range of conditions where elevated crowding has been observed, including Posterior Cortical Atrophy (Crutch & Warrington, 2007, 2009), dyslexia (Geiger & Lettvin, 1987; Atkinson, 1991; Martelli et al., 2009), and infantile nystagmus (Chung & Bedell, 1995; Pascal & Abadi, 1995). Several of these conditions report similar properties to those observed herein. For instance, crowding in Posterior Cortical Atrophy exhibits a selectivity for target-flanker similarity that matches that observed in typical peripheral vision (Yong et al., 2014). In these instances, we would expect to see the same systematic pattern of errors as in the present study. In other cases, the properties of crowding appear to differ. For instance, the spatial extent of interference zones are elongated horizontally in idiopathic infantile nystagmus, unlike the radially symmetric zones observed in typical vision and the amblyopic fovea (Tailor et al., 2021). In these cases, the pattern of response errors may differ substantially. By examining the nature of the errors produced in these distinct conditions, we can thus determine the generality of this common mechanism for crowding.

## Acknowledgements

This work was funded by the Moorfields Eye Charity (ST1411F) and the UK Medical Research Council (MR/K024817/1). Thanks to Dina Prapa from Moorfields Eye Hospital for help with participant recruitment.

## Appendices

## Appendix A

### Clinical details of children with typical vision and amblyopia

**Table A1.** Clinical details of children included in the group with typical vision (N=20). Age is reported in months. Optical correction includes cylindrical and spherical values (DS = dioptres) with the appropriate axes for each eye (L = left eye, R = right eye). logMAR acuity is also reported for each eye. Results from the TNO stereo-acuity test and are reported in seconds of arc.

Appendix A: Clinical details of children with typical vision and amblyopia
| Observer | Age<br>(months) | Gender | Refractive error | logMAR<br>acuity | TNO<br>stereo |
| --- | --- | --- | --- | --- | --- |
| T1 | 69 | M | L: Plano<br>R: Plano | L: 0.00<br>R: 0.00 | 60" |
| T2 | 78 | F | L: Plano<br>R: Plano | L: 0.00<br>R: 0.00 | 60" |
| T3 | 82 | F | L: Plano<br>R: Plano | L: 0.00<br>R: 0.00 | 60" |
| T4 | 82 | F | L: Plano<br>R: Plano | L: 0.06<br>R: 0.00 | 60" |
| T5 | 94 | M | L: Plano<br>R: Plano | L: 0.00<br>R: 0.00 | 60" |
| T6 | 88 | M | L: Plano<br>R: Plano | L: 0.00<br>R: 0.00 | 60" |
| T7 | 73 | M | L: Plano<br>R: Plano | L: 0.10<br>R: 0.04 | 120" |
| T8 | 60 | M | L: Plano<br>R: Plano | L: 0.10<br>R: 0.04 | 60" |
| T9 | 54 | M | L: +0.50 DS<br>R: +1.00 DS | L: 0.16<br>R: 0.18 | 120" |
| T10 | 75 | M | L: +0.50 DS<br>R: +1.00 DS | L: 0.10<br>R: 0.20 | 60" |
| T11 | 84 | M | L: +0.25 DS<br>R: +0.25 DS | L: 0.00<br>R: 0.00 | 60" |
| T12 | 36 | F | L: Plano<br>R: Plano | L: 0.00<br>R: 0.10 | 60" |
| T13 | 93 | M | L: -0.75/-1.00x50°<br>R: -3.00 DS | L: 0.12<br>R: 0.20 | 120" |
| T14 | 70 | M | L: Plano<br>R: Plano | L: -0.10<br>R: -0.10 | 60" |
| T15 | 73 | M | L: Plano<br>R: Plano | L: 0.00<br>R: 0.00 | 60" |
| T16 | 73 | M | L: Plano<br>R: Plano | L: 0.00<br>R: 0.00 | 60" |
| T17 | 89 | M | L: Plano<br>R: Plano | L: 0.00<br>R: 0.00 | 60" |
| T18 | 61 | F | L: Plano<br>R: Plano | L: -0.10<br>R: -0.10 | 60" |
| T19 | 74 | F | L: Plano<br>R: Plano | L: 0.02<br>R: 0.10 | 60" |
| T20 | 56 | F | L: Plano<br>R: Plano | L: 0.00<br>R: 0.00 | 60" |

**Table A2.** Clinical details of children with strabismic amblyopia included in the amblyopic group (N=20). The “ocular alignment” column indicates the outcome of near (n) and distance (d) prism tests. ET = esotropia, XT = exotropia, L/R = hypotropia. The degree of deviation is shown in prism dioptres (^Δ^), with the amblyopic eye denoted as L or R, and prism cover test results shown for both near (n) and distance (d) when required. Remaining columns are in the same format as Table A1 above.

| Observer | Age<br>(months) | Gender | Ocular<br>alignment | Refractive error | logMAR<br>acuity | TNO<br>stereo. |
| --- | --- | --- | --- | --- | --- | --- |
| A1 | 89 | M | L ET 20Δ | L: +5.25 DS<br>R: +1.75 DS | L: 0.32<br>R: -0.06 | Nil |
| A2 | 63 | M | L ET 30Δ | L: +6.5/-1.25 ×175$^\circ$<br>R: +4.5/-0.5 ×5$^\circ$ | L: 0.36<br>R: 0.1 | Nil |
| A3 | 100 | F | R ET 40Δ | L: +2.25/-0.25 ×180$^\circ$<br>R: +1.25/-2.00 ×15$^\circ$ | L: 0.04<br>R: 0.32 | Nil |
| A4 | 62 | M | R ET 35Δ | L: +2.00 DS<br>R: +2.50/-5.00 ×180$^\circ$ | L: 0.02<br>R: 0.22 | Nil |
| A5 | 67 | F | L ET 6Δ | L: +6.25/-1.00 ×180$^\circ$<br>R: +4.25/-0.25 ×180$^\circ$ | L: 0.46<br>R: 0.00 | Nil |
| A6 | 55 | M | L ET 18Δ<br>L/R 9Δ | L: +2.00/-1.00 ×180$^\circ$<br>R: +0.50/-0.50 ×180$^\circ$ | L: 0.36<br>R: 0.00 | Nil |
| A7 | 70 | F | L ET 4Δ | L: +7.00/-0.50 ×180$^\circ$<br>R: +6.75 DS | L: 0.76<br>R: 0.00 | Nil |
| A8 | 37 | F | R ET 30Δ | L: +7.25/-1.50 ×5$^\circ$<br>R: +8.00/-1.50 ×70$^\circ$ | L: 0.12<br>R: 1.10 | Nil |
| A9 | 89 | F | n: R ET 25Δ<br>d: R ET 18$^\Delta$ | L: +7.25/-2.00 ×170$^\circ$<br>R: +7.75/-1.00 ×20$^\circ$ | L: 0.00<br>R: 0.24 | Nil |
| A10 | 76 | F | n: R ET 16$^\Delta$<br>d: R ET 10$^\Delta$ | L: +4.00/ +0.5 ×180$^\circ$<br>R: +4.50/ -0.5 ×175$^\circ$ | L: 0.00<br>R: 0.38 | Nil |
| A11 | 77 | M | L ET 35$^\Delta$ | L: +5.50/-0.50 ×180$^\circ$<br>R: +3.00/-0.50 ×180$^\circ$ | L: 0.50<br>R: 0.00 | Nil |
| A12 | 86 | F | n: R ET 10$^\Delta$<br>d: R ET 4$^\Delta$ | L: +3.00/-0.50 ×180$^\circ$<br>R: +5.00/-1.00 ×180$^\circ$ | L: 0.10<br>R: 0.36 | Nil |
| A13 | 64 | F | L XT 20$^\Delta$ | L: +6.25/-1.25 ×180$^\circ$<br>R: Plano | L: 0.50<br>R: 0.00 | Nil |
| A14 | 79 | M | R ET 8$^\Delta$ | L: 0.50/-0.25 ×180$^\circ$<br>R: 5.50/-0.75 ×180$^\circ$ | L: 0.00<br>R: 0.54 | 480" |
| A15 | 83 | M | L ET 45$^\Delta$ | L: +6.50/ -1.00 ×180$^\circ$<br>R: +4.75/ -1.25 ×175$^\circ$ | L: 0.92<br>R: 0.00 | Nil |
| A16 | 70 | F | L XT 10$^\Delta$ | L: +6.00 DS<br>R: +6.00 DS | L: 0.22<br>R: 0.00 | Nil |
| A17 | 85 | M | L XT 25$^\Delta$ | L: +7.00/-1.00 ×180$^\circ$<br>R: +1.25/-0.25 ×180$^\circ$ | L: 0.86<br>R: 0.00 | Nil |
| A18 | 73 | M | L ET 40$^\Delta$ | L: +8.00/-1.50 ×70$^\circ$<br>R: +7.50/ -1.00 ×20$^\circ$ | L: 0.72<br>R: 0.18 | Nil |
| A19 | 45 | F | L ET 10$^\Delta$ | L: +7.50/-1.00 ×70$^\circ$<br>R: +6.50/-0.50 ×180$^\circ$ | L: 0.36<br>R: 0.12 | Nil |
| A20 | 44 | M | L ET 20$^\Delta$ | L: +7.00/-0.5 ×20$^\circ$<br>R: +6.00/-1.00 ×120$^\circ$ | L: 0.44<br>R: 0.02 | Nil |

## Appendix B

### Noise model simulations of group response-error distributions

As described in the main text, a noise model was fit to the data in order to examine the response errors that would arise if crowding does not have a systematic effect on target appearance, but rather distorts or adds noise to the target orientation. Figure A1 shows the result of 1000 iterations of the best-fitting noise model (see Table A3 for parameters), fit to group data. For adults, we depict here only the extreme eccentricities, with data for 2.5° in panel A and for 15° in panel B. For unflanked targets, the model again captures the distribution of response errors well, given that the first stage of the model is identical to that of the pooling model. However, with 30° target-flanker differences, the responses clearly shift away from the target towards the flanker, whereas the simulated responses of the noise model remain centred on 0°. Similarly, although the noise model can simulate the peak of responses around the target with 90° target-flanker differences, it fails to account for the secondary peak around the flanker orientation. This is even more apparent at 15° eccentricity where the peak around the flanker orientation is higher than that around the target. Intermediate eccentricities show similar effects. The model similarly fails to account for these systematic errors in both children with typical vision (panel C) and those with amblyopia (panel D). An increase in the degree of random responses alone is clearly insufficient to account for these response-error distributions.

**Table A3.** Best-fitting parameters and Akaïke Information Criterion (AIC) values for the two-parameter noise models fit to group data.

|  | **Adult periphery** |  |  |  | **Children** |  |
| --- | --- | --- | --- | --- | --- | --- |
|  | **$2.5^\circ$** | **$5^\circ$** | **$10^\circ$** | **$15^\circ$** | **Typical** | **Amblyopic** |
| Early noise | 0.21 | 0.30 | 0.31 | 0.30 | 0.71 | 0.74 |
| Late noise | 0.41 | 1.03 | 0.98 | 1.47 | 0.90 | 1.20 |
| AIC | -254.43 | -373.13 | -280.64 | -263.21 | -391.37 | -420.30 |

**Figure A1.**
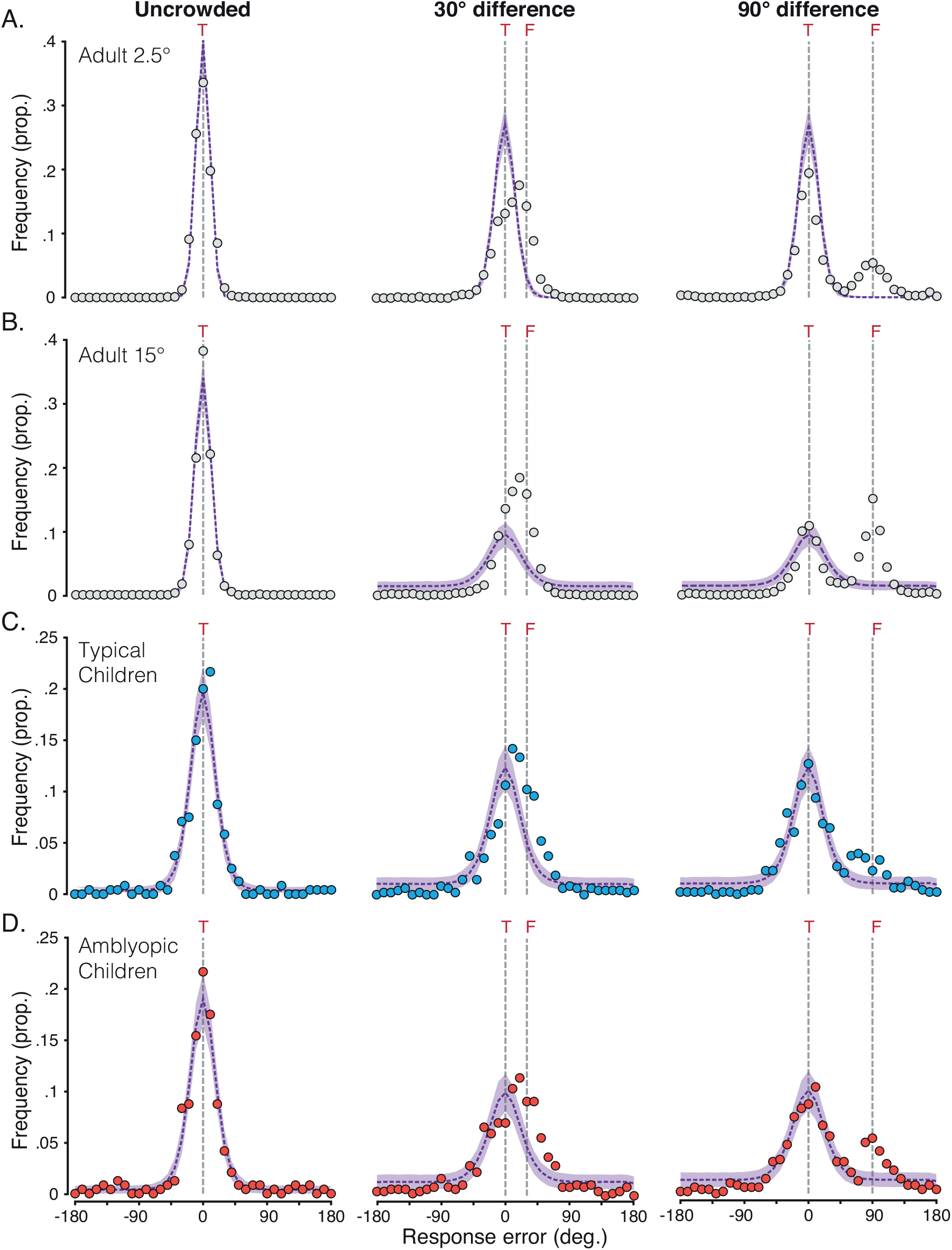
Distributions of mean response error and the simulated responses of the noise model. **A**. Response error distributions at 2.5° eccentricity, with mean values presented as light-grey dots. The purple dashed line plots the mean distribution of the noise model, with shaded areas plotting the 95% range of simulated distributions for 1000 model iterations. Dashed grey lines indicate the target location (‘T’), and for the conditions in which flankers were present, the flanker location (‘F’). **B**. Response-error distributions at 15° eccentricity and the best-fitting noise model, plotted as in panel A. **C**. Response-error distributions for children with typical vision, with mean values plotted in blue against the noise model. **D**. Response-error distributions for children with amblyopia, with mean values plotted in red against the noise model.

## Appendix C

### Pooling model simulations with single vs. multiple flanker weights

The pooling model reported in the main text used two free parameters to independently set the flanker weights in the ±30° and ±90° target-flanker difference conditions. Though this is in line with the variations in flanker weights used in some pooling models (Greenwood & Parsons, 2020), others have been successful with flanker weights that are fixed regardless of target-flanker feature differences (van den Berg, Roerdink, & Cornelissen, 2010; Harrison & Bex, 2015). Here we examine the necessity of these independent parameters for the ±30° and ±90° conditions. An alternative form of the population pooling model was fit to the group response errors, which used the same flanker-weight parameter for the two flanker conditions. The same equations were used as in the main text, with the sole change being that 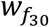 was always equal to 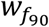. This gave three free parameters (including early and late noise), which were fit across the set of conditions in the same way as in the main text.

Figure A2 plots the mean of the 1000 iterations of the best-fitting three-parameter pooling model (dashed blue line), fit to group data and plotted against the best-fitting four-parameter model (solid black line, as described in the main text). As in Figure A1 of Appendix B, for adults we depict only the extreme eccentricities, with data for 2.5° in panel A and for 15° in panel B. Children with typical vision and those with amblyopia are shown in panels C and D. Best-fitting parameters for the four-parameter model are also presented in Table A4, for comparison with those of the three-parameter model in Table A5. For all groups, early noise values were highly similar to those of the four-parameter model. This can be seen in the histograms with unflanked targets, where the model fits are largely indistinguishable and follow the distribution of response errors closely.

To reproduce errors in the flanked conditions, best-fitting flanker weights for the three-parameter model fell closest to the weights used in the ±90° conditions for the four-parameter model (which were lower than those with ±30° flankers). In the three-parameter model, these lower weights were less effective at capturing the shift in the response-error distribution for the ±30° condition, with the late noise parameter tending to increase as a result in order to flatten out these distributions. As a result, the model outputs shown in Figure A2 reveal that the three-parameter simulation gives distributions that peak closer to the target than the peak of the response-error distributions made by observers. The four-parameter model is clearly better able to capture this shift for each group. Although the flanker weight parameters in the ±90° condition are similar for the two models, the higher late noise in the 3-parameter model tends to flatten out the error distributions, underestimating the target peak in particular. Again, the four-parameter model provides better fits to the data.

Altogether, the three-parameter model does capture many of the key properties of the errors induced by crowding – the shift in the response-error distributions to orientations between the target and flanker values in the ±30° condition, as well as the bimodal error distribution in the ±90° condition. The three-parameter model also produced AIC values that were lower than those of the two-parameter noise model (which indicates better fits with correction for the number of parameters; compare Table A3). However, AIC values were lower for the four-parameter model than the three-parameter version for all groups (see Tables A4 and A5). The variation in flanker weights with target-flanker similarity clearly gives some benefit to the performance of the model.

**Table A4.** Best-fitting parameters and Akaïke Information Criterion (AIC) values for the four-parameter pooling models fit to group data (corresponding to those reported in the main text). 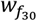 corresponds to the flanker weight for the ±30° flanker condition, with 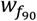 used for the ±90° condition.

|  | Adult periphery |  |  |  | Children |  |
| --- | --- | --- | --- | --- | --- | --- |
|  | 2.5° | 5° | 10° | 15° | Typical | Amblyopic |
| Early noise | 0.31 | 0.32 | 0.26 | 0.29 | 0.71 | 0.62 |
| Late noise | 1.19 | 1.19 | 0.76 | 0.73 | 1.38 | 1.85 |
| $w_{f_{30}}$ | 0.50 | 0.50 | 0.62 | 0.64 | 0.54 | 0.57 |
| $w_{f_{90}}$ | 0.35 | 0.42 | 0.50 | 0.52 | 0.32 | 0.36 |
| AIC | -502.42 | -523.42 | -560.65 | -471.00 | -490.31 | -510.42 |

**Table A5.**
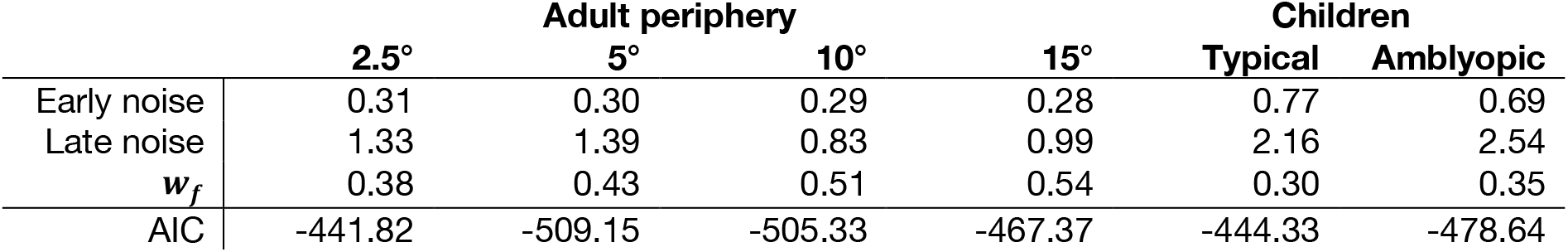
Best-fitting parameters and Akaïke Information Criterion (AIC) values for the three-parameter pooling models fit to group data (as discussed in Appendix C). *w*_*f*_ corresponds to the flanker weights, which were identical for the two flanked conditions.

|  | Adult periphery |  |  |  | Children |  |
| --- | --- | --- | --- | --- | --- | --- |
|  | 2.5° | 5° | 10° | 15° | Typical | Amblyopic |
| Early noise | 0.31 | 0.30 | 0.29 | 0.28 | 0.77 | 0.69 |
| Late noise | 1.33 | 1.39 | 0.83 | 0.99 | 2.16 | 2.54 |
| $w_f$ | 0.38 | 0.43 | 0.51 | 0.54 | 0.30 | 0.35 |
| AIC | -441.82 | -509.15 | -505.33 | -467.37 | -444.33 | -478.64 |

**Figure A2.**
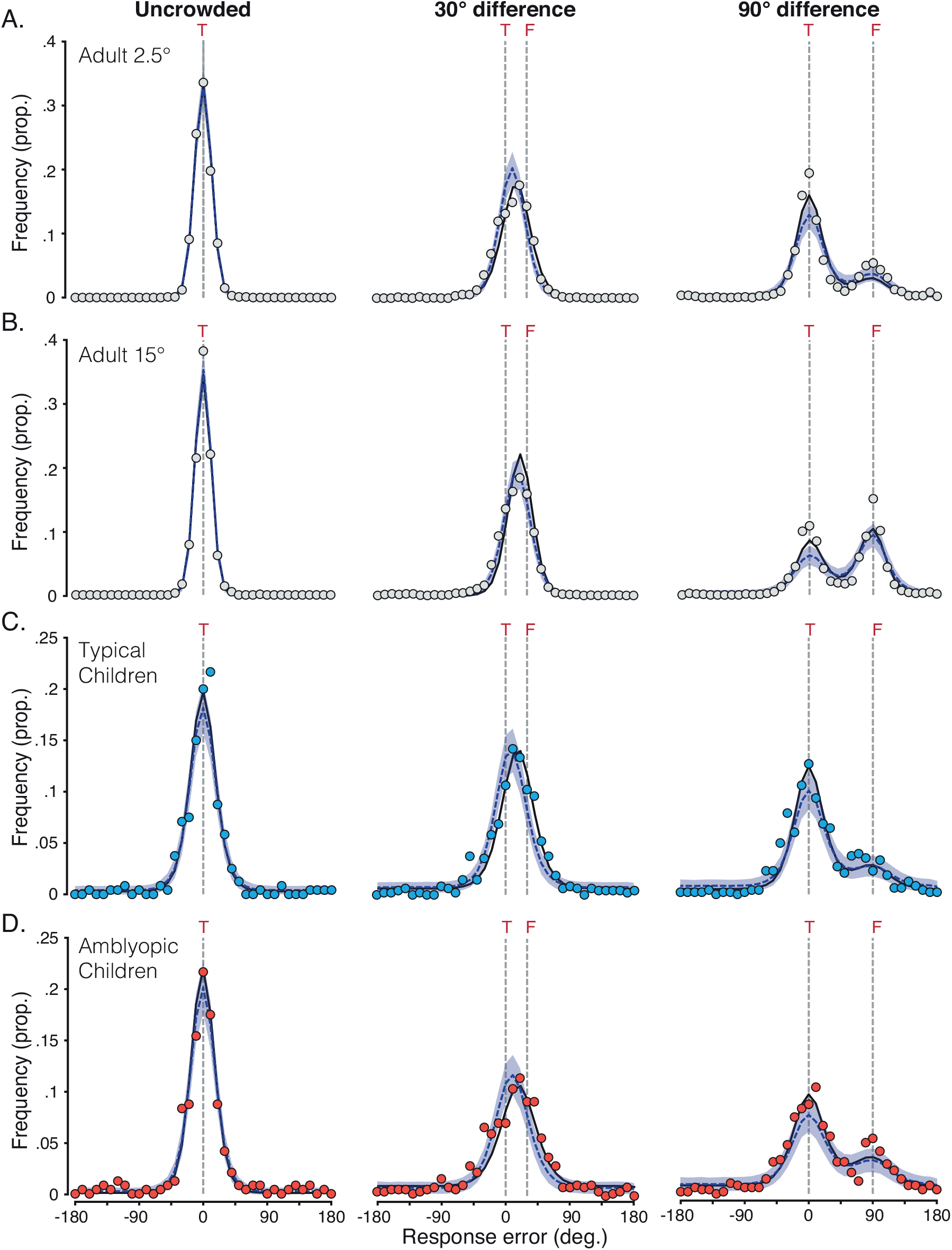
Distributions of mean response error and the best-fitting pooling model with 3 vs. 4 free parameters. **A**. Response error distributions at 2.5° eccentricity, with mean values presented as light-grey dots. The dark-blue dashed line plots the mean distribution of the population pooling model with 3 free parameters (using a single weight for both flanker conditions), with shaded areas plotting the 95% range of simulated distributions for 1000 model iterations. The solid black line shows the mean distribution of the population pooling model with 4 free parameters (as in Figures 4 and 5, here without error for clarity). Dashed grey lines indicate the target location (‘T’), and where present, the flanker location (‘F’). **B**. Response-error distributions at 15° eccentricity and the best-fitting pooling models, plotted as in panel A. **C**. Response-error distributions for children with typical vision, with mean values plotted in blue against the pooling models. **D**. Response-error distributions for children with amblyopia, with mean values plotted in red against the pooling models.

## Appendix D

### Best-fitting simulations of the models for individual data

Both pooling and noise models were also fit to individual data, as outlined in the main text. Here we plot examples to illustrate variations in the success of these models. Figure A3 plots the responses of four adult observers. Panels A and B plot data from 2.5° eccentricity. The first observer (S4) exhibited a large difference in the AIC for the two models. Although both perform equivalently for the unflanked condition (given the identical first stage of the models), their divergence is clear in the flanked conditions. In the ±30° condition, the pooling model successfully captures the shift of the response-error distribution towards the flanker, while the noise model remains centred on 0°. The secondary peak at 90° is also missed by the noise model, as in the fits to group data. In contrast, observer S5 (panel B) exhibited a lower AIC value for the noise model at this eccentricity. Although the response error distribution in the ±30° condition is lower and broader than that of the unflanked condition, it fails to show a shift towards the flanker orientation of a magnitude equivalent to that seen in other observers and the group data. Consequently, although the slightly shifted distribution of the pooling model captures this pattern better than the noise model, this improvement is not significantly greater than the fit obtained by the noise model. This is similarly true in the ±90° condition where the secondary peak in response errors is absent. The models are thus difficult to distinguish for this observer given the lower rate of errors in the flanked conditions relative to that seen in other observers.

Similar patterns can be seen for the 15° eccentricity data for two example observers in panels C and D. Observer S2 (panel C) exhibited a large difference in the AIC for the two models, driven by a clear pattern of systematic errors that is well captured by the pooling model. Contrast this with observer S1 (panel D) who showed one of the smallest AIC differences at this eccentricity. Systematic errors are less evident here, making the fit between the models harder to distinguish. Nonetheless, the shift at 30° in particular was sufficient to produce a lower AIC value for the pooling model than the noise model, indicative of a better fit.

Example fits to data from individual children are shown in Figure A4. Panels A and B plot example children with typical vision. The first (T6) had one of the largest AIC differences between the models, driven by the clear shift in their responses in the ±30° condition and the secondary peak in their responses in the ±90° condition. The noise model provides a poor account of these errors given that its distributions remain centred on 0°. Observer T2 however shows a lower degree of response errors, particularly in the 90° condition, making the models more difficult to distinguish. Nonetheless, the small shift in the response-error distribution of the 30° condition is sufficient to give a lower AIC value for the pooling model in this individual (and indeed in all children with typical vision).

Panels C and D show example individuals from the amblyopic group. Observer A11 (panel C) again shows a clear shift in response errors in the ±30° condition and a secondary peak in their responses in the ±90° condition, which the pooling model is well suited to describe. As above, the noise model provides a poor account of these errors given that its distributions remain centred on 0°. Observer A8 (panel D) differs in the sheer noisiness of their responses. Here there is clearly a distribution centred around the target, though there is little evidence for any change in these responses when flankers were added. As a result, the pooling and noise models provide equivalent fits to the data, and the penalisation of the pooling model for its greater number of parameters results in lower AIC values for the noise model. The failure of the pooling model in this case is thus driven by the noisy response-error distribution. This pattern was evident in 2 of the 20 observers in the amblyopic group, though the remainder had response error distributions that were more clearly systematic and thus better described by the pooling model.

**Figure A3.**
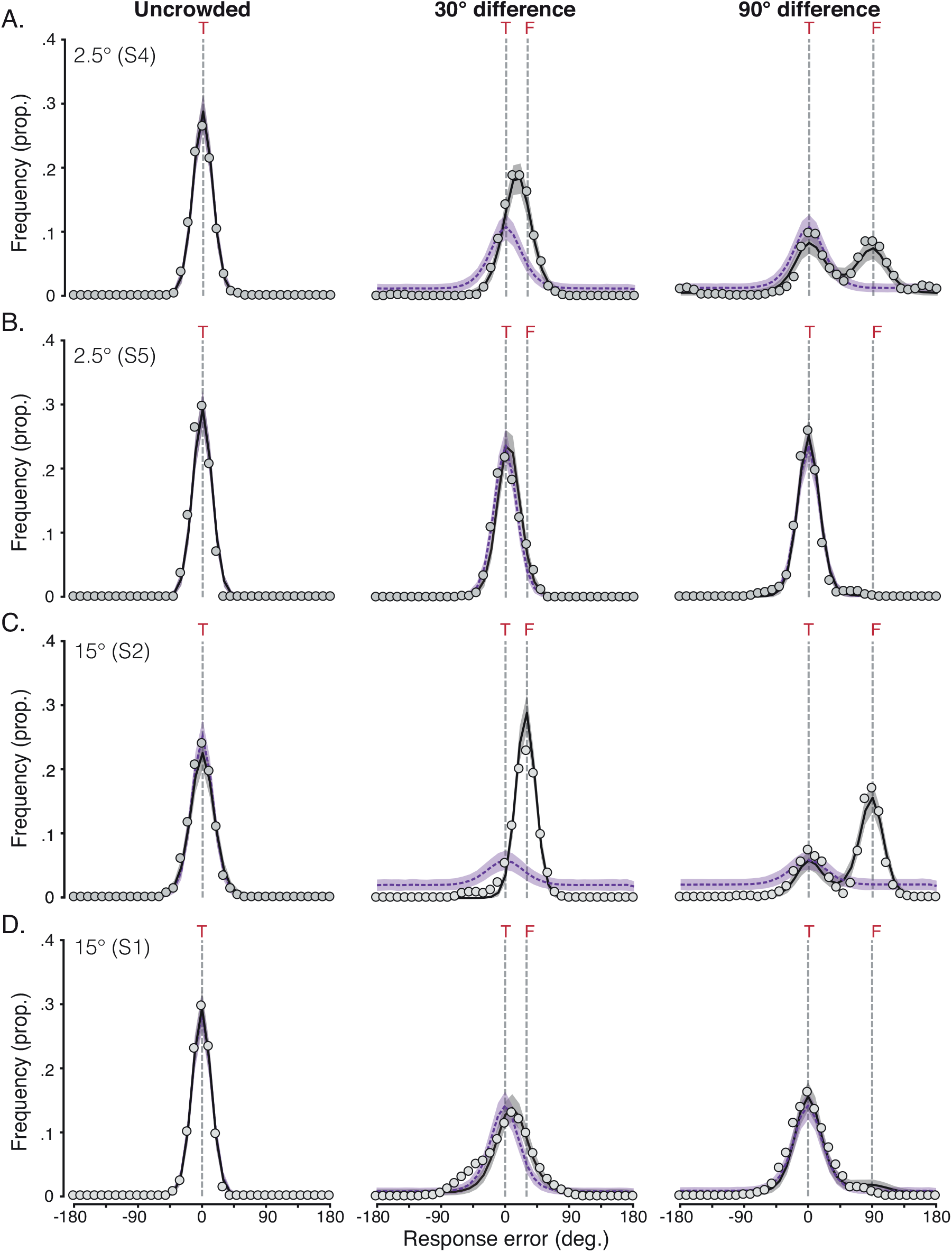
Individual data from the adult periphery and corresponding fits of the pooling and noise models. **A**. Response error distributions for one individual (S4) at 2.5° eccentricity, with mean values presented as light-grey dots. The black solid line plots the mean distribution of the pooling model, with the purple dashed line plotting the noise model. Shaded areas plot the 95% range of simulated distributions. Dashed grey lines indicate the target location (‘T’), and for the conditions in which flankers were present, the flanker location (‘F’). **B**. Response error distributions for another individual (S5) at 2.5° eccentricity and associated model fits, plotted as in panel A. **C-D**. Response error distributions for individuals S2 (C) and S1 (D) at 15° eccentricity and associated model fits, plotted as in panel A.

**Figure A4.**
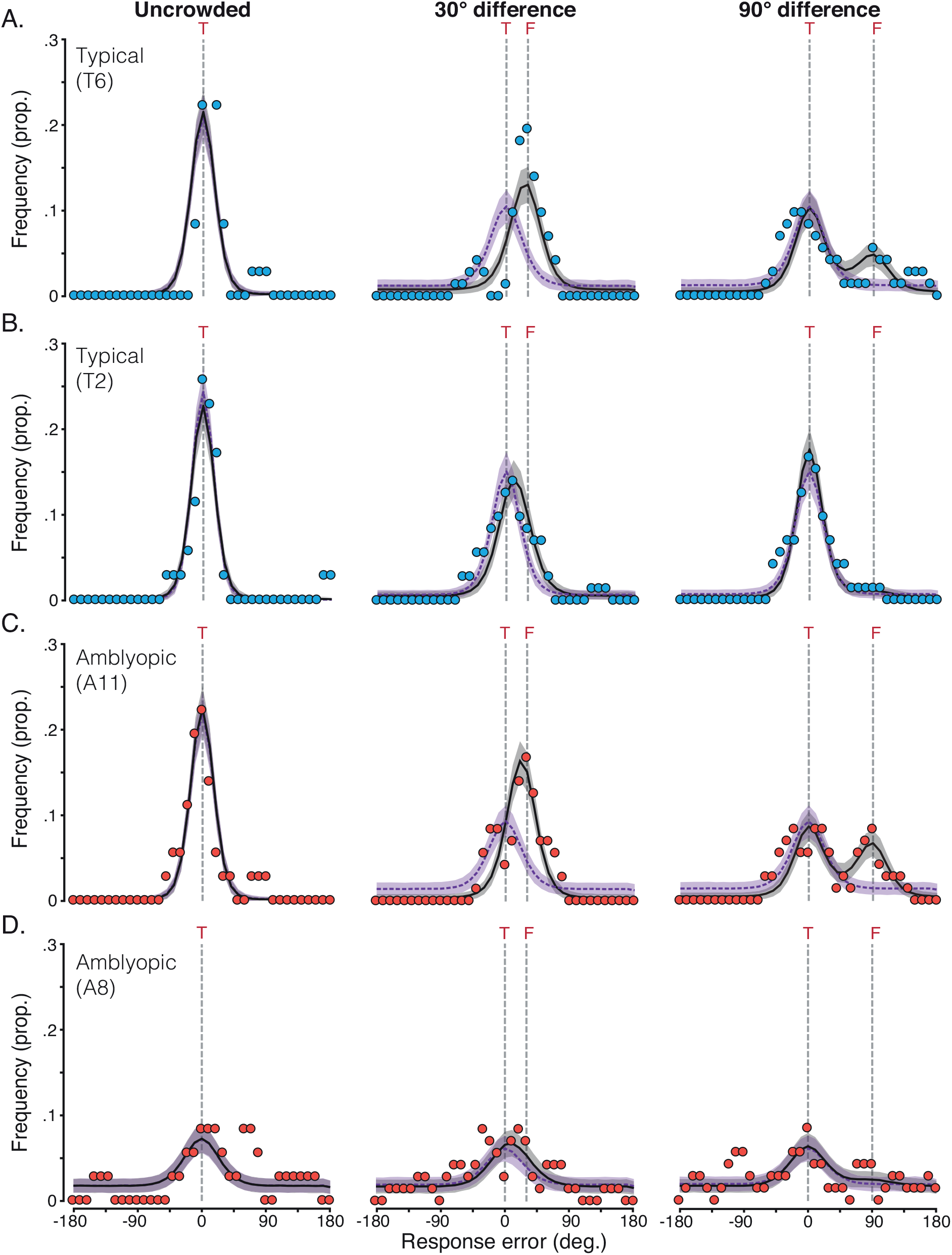
Individual data from children and corresponding fits of the pooling and noise models. **A**. Response error distributions for one child with typical vision (T6), with mean values presented as blue dots. The black solid line plots the mean distribution of the pooling model, with the purple dashed line plotting the noise model. Shaded areas plot the 95% range of simulated distributions. Dashed grey lines indicate the target location (‘T’), and for the conditions in which flankers were present, the flanker location (‘F’). **B**. Response error distributions for another child with typical vision (T2) and associated model fits, plotted as in panel A. **C-D**. Response error distributions for two children with amblyopia – A11 (C) and A8 (D) and associated model fits, plotted as in panel A.

### Footnotes

i Code available at http://github.com/eccentricvision/KidsCrowdModels

